# Cross-Stream Self-Supervised Learning Inspired by Feature-Biased Visual Processing in the Brain

**DOI:** 10.64898/2026.11.09.765087

**Authors:** Zhixian Han, Yi-Fan Li, Elizabeth M. Edwards, Anne B. Sereno

## Abstract

Humans and other animals develop rich visual representations without explicit object-category labels. Self-supervised learning (SSL) offers a computational framework for studying how such representations emerge from visual experience. Many contrastive SSL models, such as SimCLR, learn by being robust to augmented views of the same image, which may fail to preserve visual attributes such as color and orientation. Inspired by feature-specific visual processing streams in the brain, we propose BioCLR, an SSL framework that uses color- and orientation-biased networks to guide representation learning. We combine Bio-CLR with SimCLR into a BioCLR hybrid model, pretrain it on ImageNet-100, and evaluate it on this dataset and four transfer datasets. In frozen linear evaluation, the BioCLR hybrid outperforms SimCLR across all datasets and mitigates negative transfer on a handwritten-character dataset. In limited-label fine-tuning, it outperforms SimCLR in most transfer conditions. Compared with the AugSelf and RotNet hybrids, which incorporate color and orientation information into SimCLR in other ways, the BioCLR hybrid shows the best overall performance, particularly in limited-label fine-tuning. Ablation analyses suggest that orientation guidance helps more than color guidance on these datasets, and that combining both generally improves frozen linear evaluation performance. Together, these findings suggest that incorporating feature-specific information into SSL can improve representation quality and model adaptability, and that the learning mechanism used to incorporate this information matters. For neuroscience, these results suggest that specialized visual streams in the brain may provide learning signals that guide visual representation learning without explicit supervision. For machine learning, they imply that principles of feature-biased processing in visual cortex may improve SSL.

## 1 Introduction

Humans and other animals develop rich visual representations from natural visual experience without requiring explicit category labels for every object they encounter. In humans, object-category representations develop in the ventral visual cortex during infancy without explicit labels accompanying each visual experience (O’Doherty et al., 2026). Similarly, face-selective areas develop as infant macaques gain visual experience with faces, without requiring explicit labels identifying those faces (Livingstone et al., 2017). In contrast, macaques raised without seeing faces fail to develop normal face-selective areas, suggesting that visual experience with faces is essential for the development of face-selectivity (Arcaro et al., 2017). This raises an important question for both neuroscience and machine learning: what mechanisms allow the visual system to develop useful object representations under limited external supervision?

One possibility is that the visual system can learn object representations without external supervision by learning and becoming robust to different views of the same object. For example, changes in illumination alter the light reflected from an object while its identity remains unchanged (Hurlbert, 2019; Pastilha & Hurlbert, 2022). Similarly, movements of objects and observers allow the same object to be experienced from different viewpoints (DiCarlo & Cox, 2007; Wallis et al., 2009). These differing experiences may allow visual representations to develop without explicit category labels by becoming robust to changes in color and viewpoint. However, becoming invariant to these changes may also cause visual representations to lose important visual attribute information. For example, color invariance could reduce color information that is useful for distinguishing an object’s identity or state, while strong viewpoint invariance could reduce information about its orientation or spatial configuration. Here, we propose an additional biologically plau-sible mechanism for visual learning: general visual representations may also develop by aligning with more specialized representations generated by different visual streams in response to the same sensory input.

Many neuroscience studies have suggested that there are two main cortical visual pathways in the brain, a ventral pathway specialized for object recognition and a dorsal pathway specialized for spatial recognition (Ungerleider & Mishkin, 1982; Mishkin et al., 1983; Felleman & Van Essen, 1991; Goodale & Milner, 1992; Han & Sereno, 2024a). In addition, visual cortex contains multiple partially specialized feature-biased systems within each main cortical pathway, including systems involved in color, orientation, form, motion, faces, bodies, and other socially relevant stimuli (Gegenfurtner, 2003; Seymour et al., 2010; Rentzeperis et al., 2014; Taylor & Xu, 2022; Murata et al., 2000; Valyear et al., 2006; Aflalo & Graziano, 2011; Kravitz et al., 2011; 2013; Pitcher & Ungerleider, 2021; Taubert et al., 2022). These systems may retain different aspects of visual attributes to produce related, but distinct, representations of the same sensory input (Sereno et al., 2020). Computational modeling studies have also suggested that different artificial visual pathways can develop distinct representations of a visual scene that support different aspects of visual recognition and localization (Rueckl et al., 1989; Jacobs et al., 1991; Scholte et al., 2018; Bakhtiari et al., 2021; Choi et al., 2023; Han & Sereno, 2022; 2023). If different pathways and systems emphasize different visual attributes to achieve different goals, then the representations they generate may provide internal constraints that help shape and constrain more general-purpose visual representations. Together, these findings motivate the hypothesis that interactions among specialized visual streams may play an important role in visual representation learning by encouraging and retaining the encoding of feature-specific information.

Self-supervised learning (SSL) provides an important computational framework for studying how visual representations in the brain can develop without explicit object-category labels. In contrastive SSL, neural networks are trained by comparing positive (match) and negative (nonmatch) pairs of inputs and minimizing a contrastive loss function (Hadsell et al., 2006; Tian et al., 2020a; Hénaff, 2020). SimCLR is a simple and effective framework for contrastive learning of visual representations (Chen et al., 2020a). During SimCLR training, two augmented views are generated from the same image and treated as a positive pair, whereas augmented views generated from different images are treated as negative pairs. The network is trained so that representations of positive pairs become similar and representations of negative pairs become dissimilar. This procedure allows SimCLR to learn visual representations without object-category labels. However, one potential limitation of SimCLR is that its objective encourages agreement between augmented views of the same image but does not actively promote the encoding or retention of potentially useful visual attribute information which may differ across views. Consequently, information about some visual attributes (e.g., color and orientation) may be weakened as the model learns to match representations of transformed inputs (Xiao et al., 2021; Lee et al., 2021).

To address this limitation in the current study, we investigate whether incorporating separate specialized (color and orientation) artificial visual streams (BioCLR) into SimCLR to create a BioCLR hybrid model (SimCLR + BioCLR) improves general-purpose visual representation learning by providing additional SSL signals from these specialized streams. BioCLR uses convolutional neural networks to model a color stream and an orientation stream. The color stream is trained to classify the hue of the average color in the central region of each image, whereas the orientation stream is trained to classify image orientation. After training, these networks are frozen and used as teacher networks to guide a student network through their specialized representations. For each input image, BioCLR encourages the student’s representation to align with the corresponding representations produced by both teachers while distinguishing it from student representations of other images. Therefore, BioCLR combines cross-stream alignment for the same sensory input with contrastive separation of representations for different inputs. When combined with SimCLR, these additional learning signals may provide specialized color and orientation constraints that help preserve useful visual attribute information in general-purpose visual representations. This teacher–student design is related to knowledge distillation (Hinton et al., 2015), particularly contrastive representation distillation (Tian et al., 2020b). To our knowledge, this is the first use of feature-biased teachers, motivated by feature-biased streams in visual cortex, to complement augmentation-based SSL. The overall BioCLR hybrid model is illustrated in Figure 1 and details are provided in Appendix A. Although the BioCLR hybrid model is designed to support general-purpose visual representation learning, our current study evaluates its learned representations only in terms of visual object recognition.

**Figure 1:**
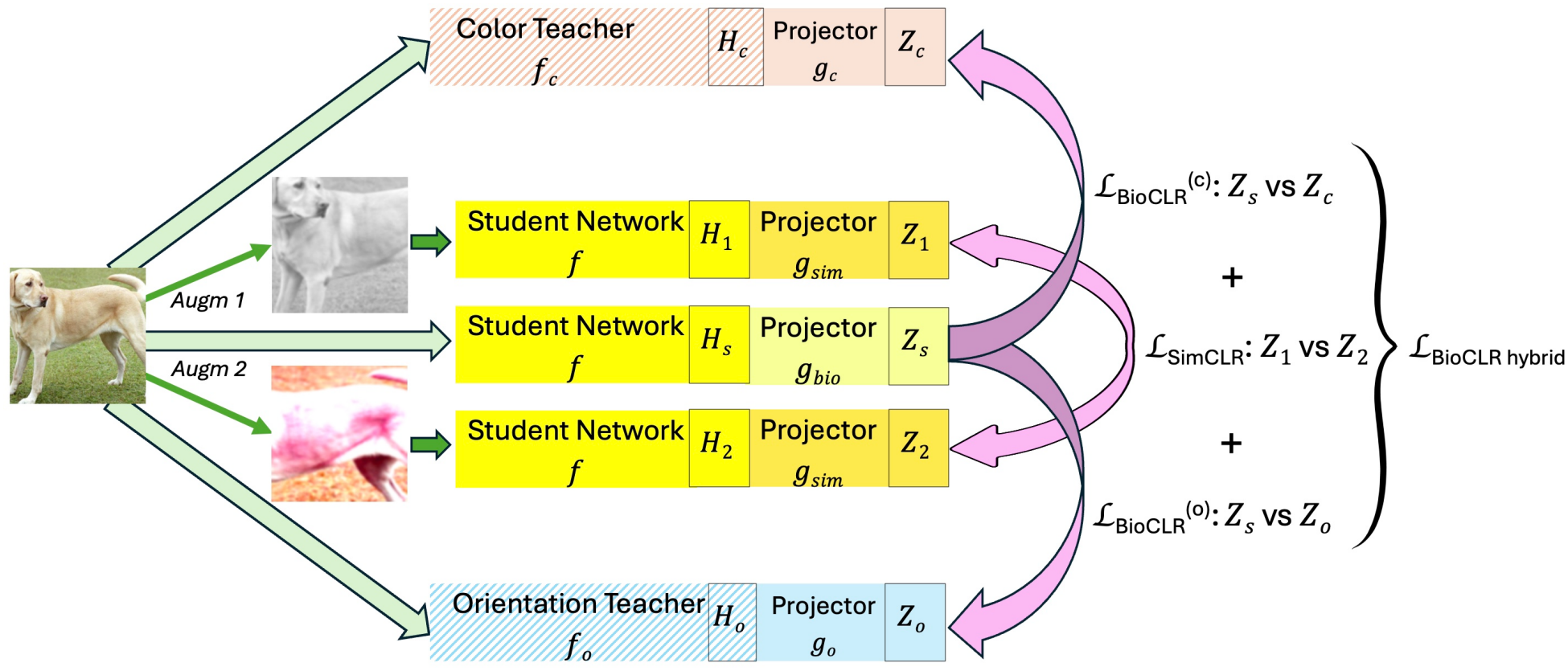
BioCLR hybrid model (SimCLR + BioCLR). Full details are provided in Appendix A. All three student networks share the weights *f*. Two augmented views of each image follow the SimCLR pathway of Figure A.6(a). The projector *g*_sim_ maps their student representations *H*_1_ and *H*_2_ to *Z*_1_ and *Z*_2_, respectively. The loss _SimCLR_ compares *Z*_1_ and *Z*_2_. The BioCLR teachers *f*_*c*_ and *f*_*o*_ were trained as in Figure A.6(b) and stay fixed (hatched). The upright original preprocessed image follows the BioCLR pathway of Figure A.6(c). The projectors *g*_*c*_ and *g*_*o*_ map its teacher representations *H*_*c*_ and *H*_*o*_ to *Z*_*c*_ and *Z*_*o*_, respectively. The projector *g*_bio_ maps its student representation *H*_*s*_ to *Z*_*s*_. The losses 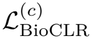 and 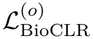 compare *Z*_*s*_ with the teacher embeddings *Z*_*c*_ and *Z*_*o*_, respectively. Each pink arrow points from an anchor representation to its positive. The total loss is 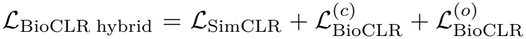. Only the student and the projectors are updated. Dog images adapted from a photo by Elf and Djmirko (Wikimedia Commons, CC BY-SA 3.0).

We address whether adding specialized streams as self-supervised teacher networks in BioCLR to SimCLR improves downstream object recognition accuracy under two evaluation settings: (1) frozen linear evaluation and (2) limited-label fine-tuning. For frozen linear evaluation, we also compare the BioCLR hybrid model with post-hoc concatenation of SimCLR and specialized teacher representations. In addition, we compare the BioCLR hybrid model with two alternative hybrid models (the AugSelf hybrid (SimCLR + AugSelf), adapted from Lee et al., 2021; and the RotNet hybrid (SimCLR + RotNet), adapted from Gidaris et al., 2018) designed to promote the encoding of color and orientation in SSL through different approaches. Finally, using an ablation study, we examine how the color- and orientation-biased teacher networks in BioCLR individually and jointly contribute to the performance of the BioCLR hybrid model.

## 2 Methods

Full details are provided in Appendix A. All encoders in this study, including the BioCLR student and teachers and the encoders of all comparison models, use the same ResNet-18 architecture (He et al., 2016).

### 2.1 Datasets

We use ImageNet-100 (Figure A.1), the 100-class subset of ImageNet (Deng et al., 2009) defined by Tian et al. (2020a), as the source natural image dataset for SSL pretraining and in-domain evaluation. We use its validation split as the held-out test set. We use four additional datasets to evaluate each model’s transfer performance across different visual recognition tasks. CIFAR-10 (Figure A.2) provides a near-domain transfer task involving natural object images (Krizhevsky & Hinton, 2009). Flowers102 (Figure A.3) provides a fine-grained flower recognition task in which color information is relevant (Nilsback & Zisserman, 2008). EMNIST Balanced (Figure A.4) provides a handwritten-character recognition task in which orientation information is relevant (Cohen et al., 2017). Finally, a directional subset of GTSRB (Figure A.5) provides a traffic-sign recognition task in which both color and orientation information are relevant (Stallkamp et al., 2011). Image preprocessing details are provided in Appendix A.2.

### 2.2 BioCLR hybrid model

The BioCLR hybrid model trains a student encoder with SimCLR and BioCLR components. The loss for the hybrid model is the sum of the losses of these components:

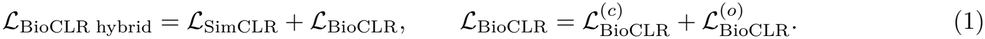

#### SimCLR component

A SimCLR projection head maps the student representations of two augmented views of image *i* to the embeddings **z**_1,*i*_ and **z**_2,*i*_. To match the number of negative pairs per anchor image in the BioCLR component loss, we modify the standard NT-Xent loss (Chen et al., 2020a) for the SimCLR component (temperature *τ*_*s*_) to use negatives only from the opposite augmented-view batch:

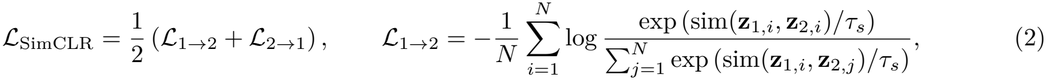

where *N* is the minibatch size, sim(*·, ·*) is cosine similarity, and *ℒ*_2*→*1_ is defined analogously.

#### BioCLR component

During teacher training only, each image is randomly rotated to one of four orientations (up, down, left, or right). The color teacher is trained to classify the hue of the average color in a centered region spanning 60% of the image height and width into one of 12 hue categories. This approximates fovea-centered color sampling. The orientation teacher is trained to classify image orientation. Both teachers are trained without human-annotated labels: the hue labels are computed from the images themselves, and the orientation labels are computed from the rotation applied to each image. After training, the teachers are frozen. The student and both teachers receive the same original preprocessed image, and separate projection heads map their representations to the embeddings **z**_*s,i*_ and **z**_*t,i*_ of image *i*, where *t* ∈ {*c, o*} denotes the color or orientation teacher. For each teacher, the BioCLR loss (temperature *τ*_*b*_) treats the student and teacher embeddings of the same image as positive pairs and the student embeddings of the other images in the minibatch as negatives:

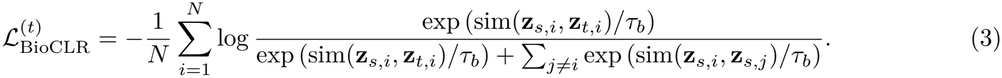

### 2.3 Comparison models

The comparison models are two baselines (untrained and supervised), standard SimCLR, a post-hoc concatenation control, two alternative hybrid models, and two ablation models. The untrained baseline is a network with randomly initialized weights. The supervised baseline is a network trained from scratch on the same labeled subsets as the pretrained models. Standard SimCLR is trained with the standard NT-Xent loss. The post-hoc concatenation control combines the representations of independently trained standard SimCLR, color-teacher, and orientation-teacher encoders. Each alternative hybrid model adds a color predictor and an orientation predictor on top of a SimCLR encoder to encourage color and orientation encoding. The predictor losses are minimized jointly with the standard NT-Xent loss. In the AugSelf hybrid, predictors are trained to predict differences in color-jitter parameters and orientation between two augmented views. In the RotNet hybrid, the SimCLR encoder also receives a randomly rotated copy of each original preprocessed image. Predictors are trained to predict its hue category and orientation. In the ablation study, the BioCLR components of the color-only and orientation-only BioCLR hybrids use only 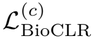 and 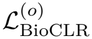 respectively.

### 2.4 Model evaluation

Only the encoder of each model (the student encoder in BioCLR hybrids) is retained and evaluated. In frozen linear evaluation, the encoder is kept fixed, and only a linear classifier is trained on the encoder representations using the full labeled training set. In limited-label fine-tuning, both the encoder and the classifier are updated using a subset of labeled training images: 1% or 10% of the training images (ImageNet-100), 0.1% or 1% of the training images (CIFAR-10 and EMNIST Balanced), 1 or 5 labeled images per class (Flowers102), and 5 or 50 labeled images per class (Directional GTSRB). Accuracy is reported for ImageNet-100, CIFAR-10, and EMNIST Balanced, whereas balanced accuracy is reported for Flowers102 and Directional GTSRB to account for differences in class frequencies. The entire experiment is repeated in three independent runs.

## 3 Results

The results are organized into five sets of analyses: teacher network evaluation (Section 3.1), frozen linear evaluation of learned representations (Section 3.2), limited-label fine-tuning (Section 3.3), a comparison of the BioCLR hybrid with the AugSelf and RotNet hybrids (Section 3.4), and an ablation study (Section 3.5). All model comparisons (Sections 3.2–3.5) are conducted on the pretraining dataset ImageNet-100 and four transfer datasets: CIFAR-10, Flowers102, EMNIST Balanced, and Directional GTSRB.

Performance of all model conditions is reported as the mean and standard deviation (mean ± SD) of test accuracy or balanced accuracy across three independent runs. Within each evaluation condition, every pair of models is compared using a two-sided Welch’s *t*-test, and the resulting *p*-values are adjusted for multiple comparisons using the Benjamini–Hochberg (BH) false discovery rate (FDR) procedure (Benjamini & Hochberg, 1995). Differences with *p*_BH_ *<* 0.05 are considered significant, and differences with 0.05 ≤ *p*_BH_ *<* 0.10 are considered marginal. Model comparisons are shown in Figures 2–7, and the corresponding tables are provided in Appendix B.

**Figure 2:**
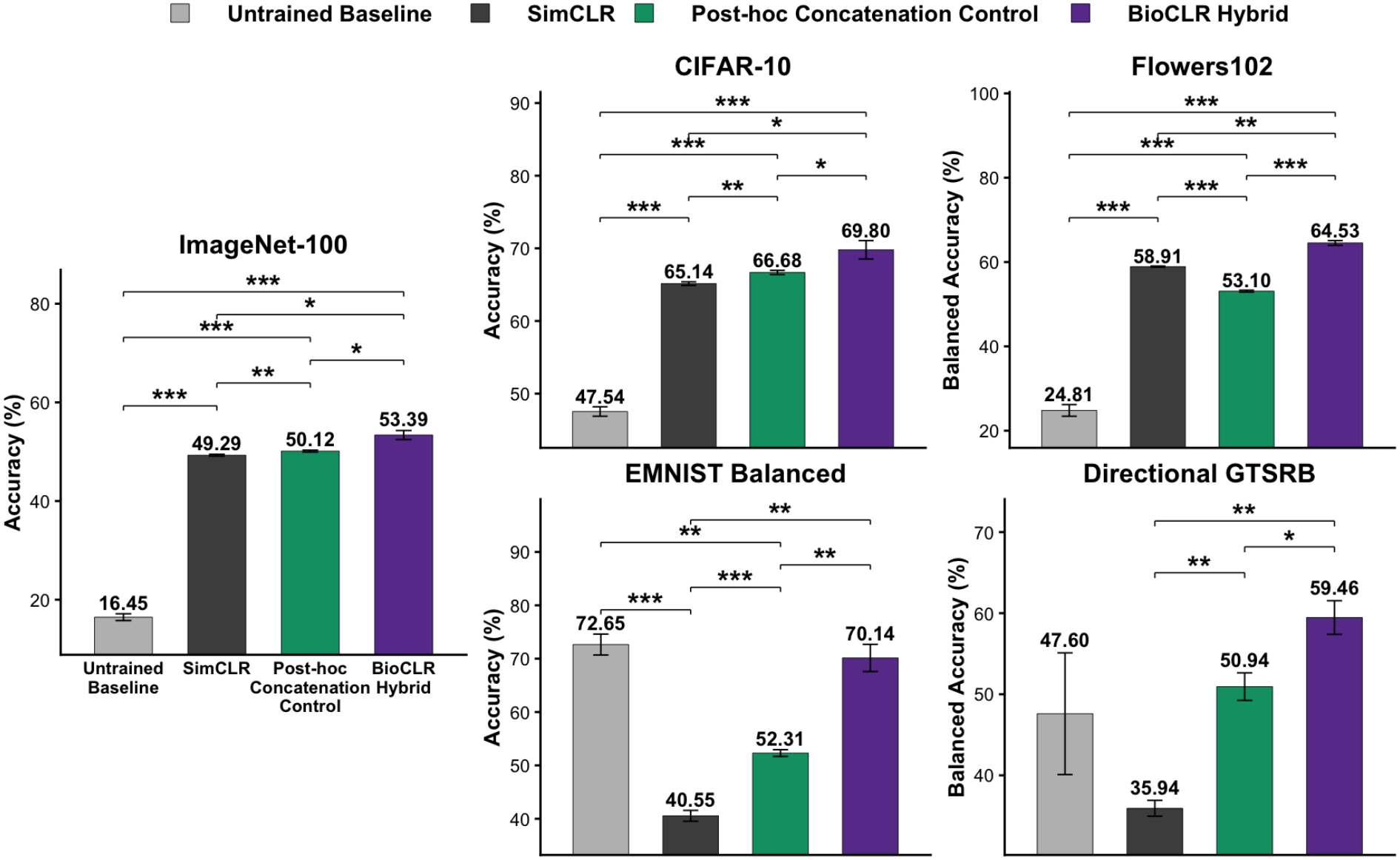
Frozen linear evaluation comparing an untrained baseline, standard SimCLR, a post-hoc concatenation control (SimCLR + color + orientation concatenation), and the BioCLR hybrid. Bars represent mean test accuracy or balanced accuracy across three independent runs and error bars show standard deviation. Statistical annotations are based on two-sided Welch’s *t*-tests with BH FDR correction across the six pairwise comparisons within each dataset. \**p*_BH_ *<* 0.05, \*\**p*_BH_ *<* 0.01, and \*\*\**p*_BH_ *<* 0.001; † indicates marginal significance (0.05 ≤ *p*_BH_ *<* 0.10).

### 3.1 Teacher Network Evaluation

On the held-out ImageNet-100 test set, the color teacher classifies the 12 image hue categories with 91.37 ± 0.19% balanced accuracy (chance: 8.33%), and the orientation teacher identifies four image orientations with 79.48 ± 0.64% accuracy (chance: 25%). Both teachers perform well above chance, suggesting that they encode color and orientation information, as expected.

### 3.2 Frozen Linear Evaluation of Learned Representations

Figure 2 and Table B.1 show the frozen linear evaluation results. The BioCLR hybrid significantly outperforms SimCLR on ImageNet-100 and on all four transfer datasets (CIFAR-10, Flowers102, EMNIST Balanced, and Directional GTSRB).

We also compare standard SimCLR and the BioCLR hybrid with an untrained baseline to assess the effects of SSL pretraining. The BioCLR hybrid significantly outperforms the untrained baseline on ImageNet-100, CIFAR-10, and Flowers102, while performing similarly to the untrained baseline on EMNIST Balanced and Directional GTSRB. In contrast, standard SimCLR performs significantly worse than the untrained baseline on EMNIST Balanced, suggesting negative transfer from ImageNet-100 pretraining to handwritten-character recognition. Similarly, on Directional GTSRB, SimCLR achieves a much lower mean than the untrained baseline, although the difference is not significant. Notably, the BioCLR hybrid significantly outperforms SimCLR on both datasets and does not show significant negative transfer relative to the untrained baseline.

In addition, we compare the BioCLR hybrid with a post-hoc concatenation control to assess whether the benefits of the hybrid model extend beyond concatenating independently learned representations. The BioCLR hybrid significantly outperforms the concatenation control on all five datasets.

### 3.3 Limited-Label Fine-Tuning

Figure 3 and Table B.2 show the limited-label fine-tuning results. On ImageNet-100 (Figure 3, top panels), the BioCLR hybrid achieves the highest mean performance with 1% labels, although its advantage over SimCLR is marginal. With 10% labels, SimCLR achieves a higher mean than the BioCLR hybrid, but this difference is also marginal. Both pretrained models significantly outperform the supervised baseline under both label conditions.

**Figure 3:**
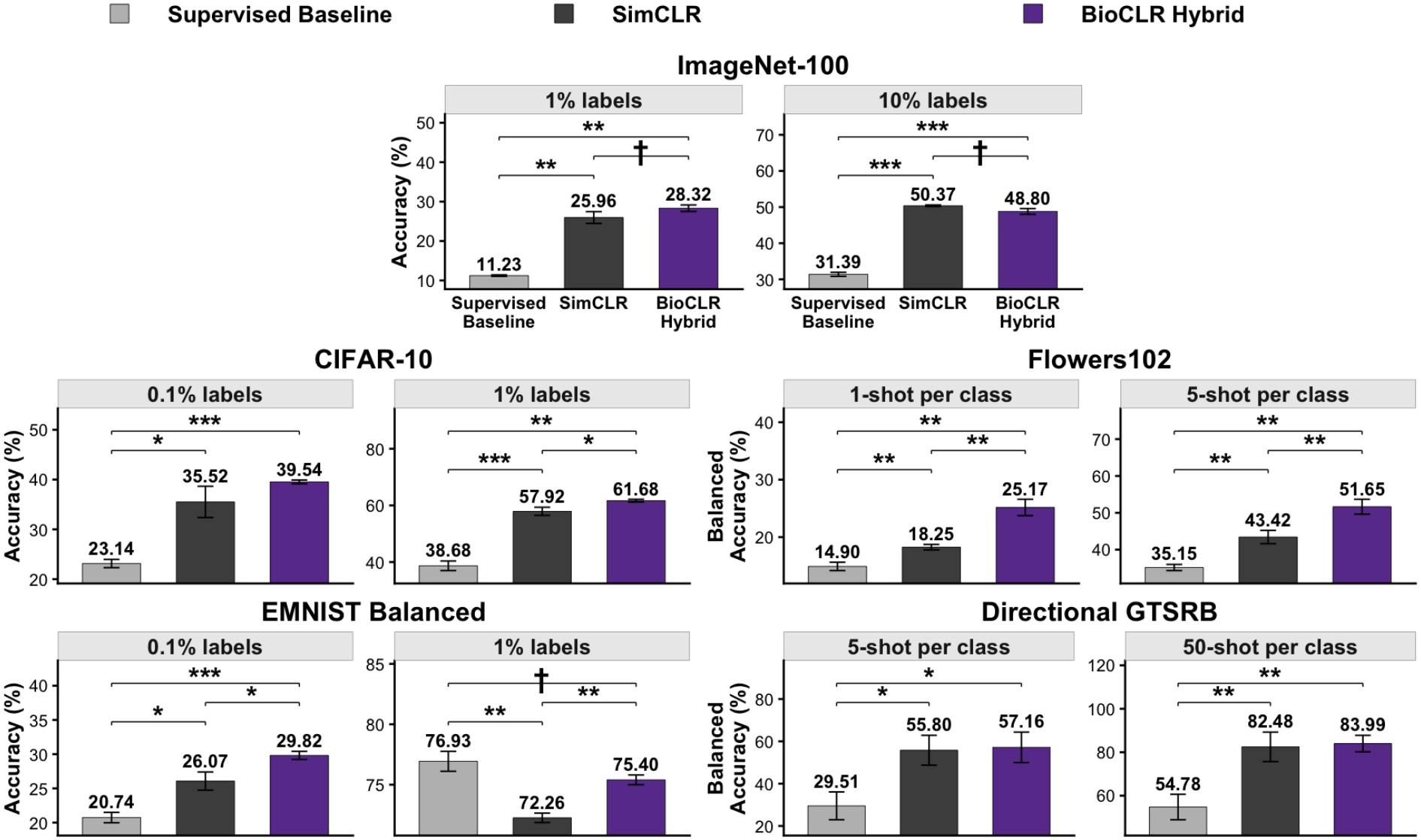
Limited-label fine-tuning comparing a supervised baseline, standard SimCLR, and the BioCLR hybrid. Bars show mean test accuracy or balanced accuracy across three independent runs and error bars show standard deviation. Statistical annotations are based on two-sided Welch’s *t*-tests with BH FDR correction across the three pairwise comparisons within each dataset and label condition. \**p*_BH_ *<* 0.05, \*\**p*_BH_ *<* 0.01, and \*\*\**p*_BH_ *<* 0.001; † indicates marginal significance (0.05 ≤ *p*_BH_ *<* 0.10).

Across the four transfer datasets, the BioCLR hybrid generally outperforms SimCLR. Specifically, the Bio-CLR hybrid significantly outperforms SimCLR on CIFAR-10 with 1% labels, Flowers102 with both 1 and 5 shots per class, and EMNIST Balanced with both 0.1% and 1% labels. On CIFAR-10 with 0.1% labels and Directional GTSRB with both 5 and 50 shots per class, the BioCLR hybrid achieves higher mean performance than SimCLR, although these differences are not statistically significant.

Compared with the supervised baseline trained from scratch, both pretrained models generally achieve higher performance across the transfer datasets, with the exception of EMNIST Balanced with 1% labels. In this condition, the supervised baseline achieves the highest mean performance, followed by the BioCLR hybrid and SimCLR. Standard SimCLR performs significantly worse than the supervised baseline, suggesting negative transfer from ImageNet-100 pretraining. In contrast, the BioCLR hybrid significantly outperforms SimCLR, although its performance remains marginally lower than that of the supervised baseline.

### 3.4 Comparison of the BioCLR Hybrid With the AugSelf and RotNet Hybrids

Figure 4 and Table B.3 show the frozen linear evaluation results. On ImageNet-100 (Figure 4, left panel), the BioCLR hybrid achieves the highest mean performance and significantly outperforms both the AugSelf and RotNet hybrids. However, this advantage does not consistently extend to the transfer datasets. On CIFAR-10, EMNIST Balanced, and Directional GTSRB, no significant differences are observed among the three models. On Flowers102, the AugSelf hybrid significantly outperforms both the BioCLR and RotNet hybrids. On Directional GTSRB, both the AugSelf and RotNet hybrids achieve higher means than the BioCLR hybrid, but the differences are marginal.

**Figure 4:**
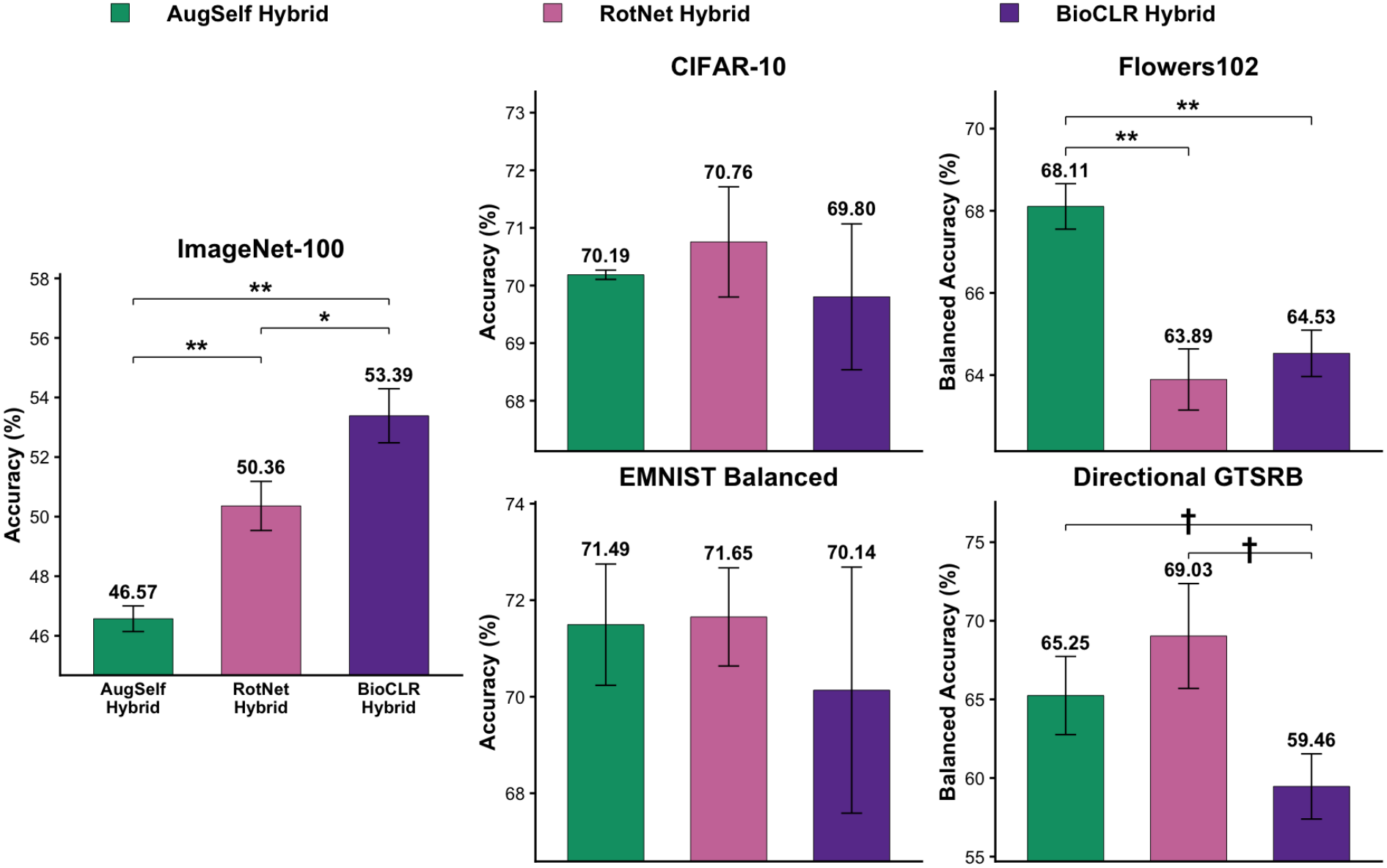
Frozen linear evaluation comparing the AugSelf, RotNet, and BioCLR hybrids. Bars show mean test accuracy or balanced accuracy across three independent runs and error bars show standard deviation. Statistical annotations are based on two-sided Welch’s *t*-tests with BH FDR correction across the three pairwise comparisons within each dataset. \**p*_BH_ *<* 0.05, \*\**p*_BH_ *<* 0.01, and \*\*\**p*_BH_ *<* 0.001; † indicates marginal significance (0.05 ≤ *p*_BH_ *<* 0.10).

Figure 5 and Table B.4 show the limited-label fine-tuning results. The BioCLR hybrid generally achieves the highest performance among the three models, with the highest mean in eight of the ten conditions. On ImageNet-100 (Figure 5, top panels), the BioCLR hybrid significantly outperforms both the AugSelf and RotNet hybrids with both 1% and 10% labels. Across the transfer datasets, the BioCLR hybrid significantly outperforms the AugSelf hybrid on CIFAR-10 with 1% labels, Flowers102 with 5 shots per class, and EMNIST Balanced with 1% labels, with a marginal advantage on Flowers102 with 1 shot per class. The BioCLR hybrid also significantly outperforms the RotNet hybrid on CIFAR-10 and EMNIST Balanced with 1% labels, with a marginal advantage on Flowers102 with 5 shots per class. However, no significant differences among the three hybrid models are observed on CIFAR-10 and EMNIST Balanced with 0.1% labels, or on Directional GTSRB with either 5 or 50 shots per class. Overall, the BioCLR hybrid significantly outperforms the AugSelf hybrid in five of the ten conditions and the RotNet hybrid in four of the ten conditions, without performing significantly worse than either alternative hybrid model in any condition.

**Figure 5:**
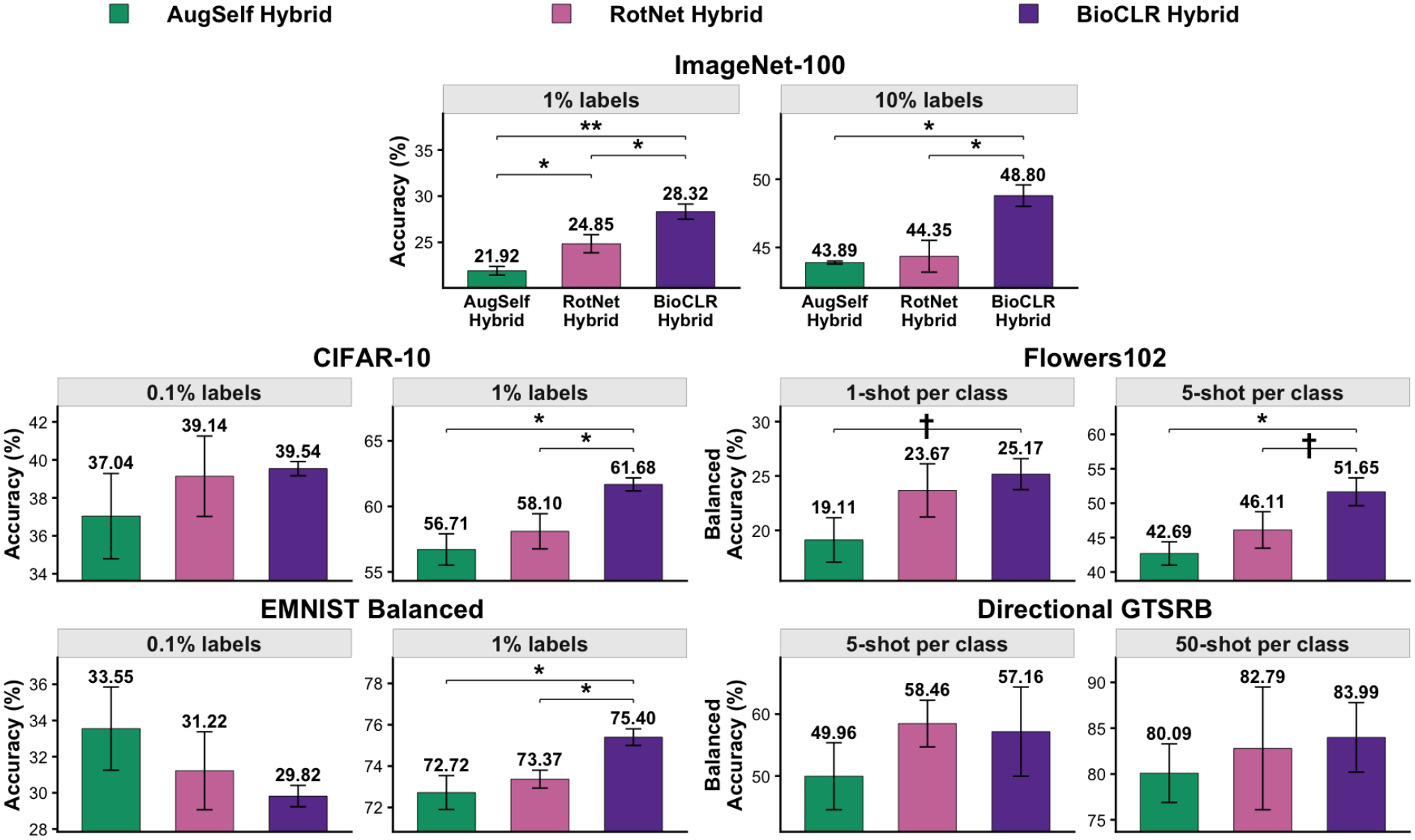
Limited-label fine-tuning comparing the AugSelf, RotNet, and BioCLR hybrids. Bars show mean test accuracy or balanced accuracy across three independent runs and error bars show standard deviation. Statistical annotations are based on two-sided Welch’s *t*-tests with BH FDR correction across the three pairwise comparisons within each dataset and label condition. \**p*_BH_ *<* 0.05, \*\**p*_BH_ *<* 0.01, and \*\*\**p*_BH_ *<* 0.001; † indicates marginal significance (0.05 ≤ *p*_BH_ *<* 0.10).

### 3.5 Ablation Study

Figure 6 and Table B.5 show the frozen linear evaluation results. The full BioCLR hybrid model achieves the highest mean performance across all five datasets and significantly outperforms both BioCLR color-only and BioCLR orientation-only hybrid models on ImageNet-100, CIFAR-10, Flowers102, and EMNIST Balanced. On Directional GTSRB, the full model achieves a marginal advantage over the color-only model, whereas its advantage over the orientation-only model is not statistically significant. Comparing the two single-teacher models, the orientation-only model significantly outperforms the color-only model on ImageNet-100, CIFAR-10, Flowers102, and EMNIST Balanced, with a marginal advantage on Directional GTSRB.

**Figure 6:**
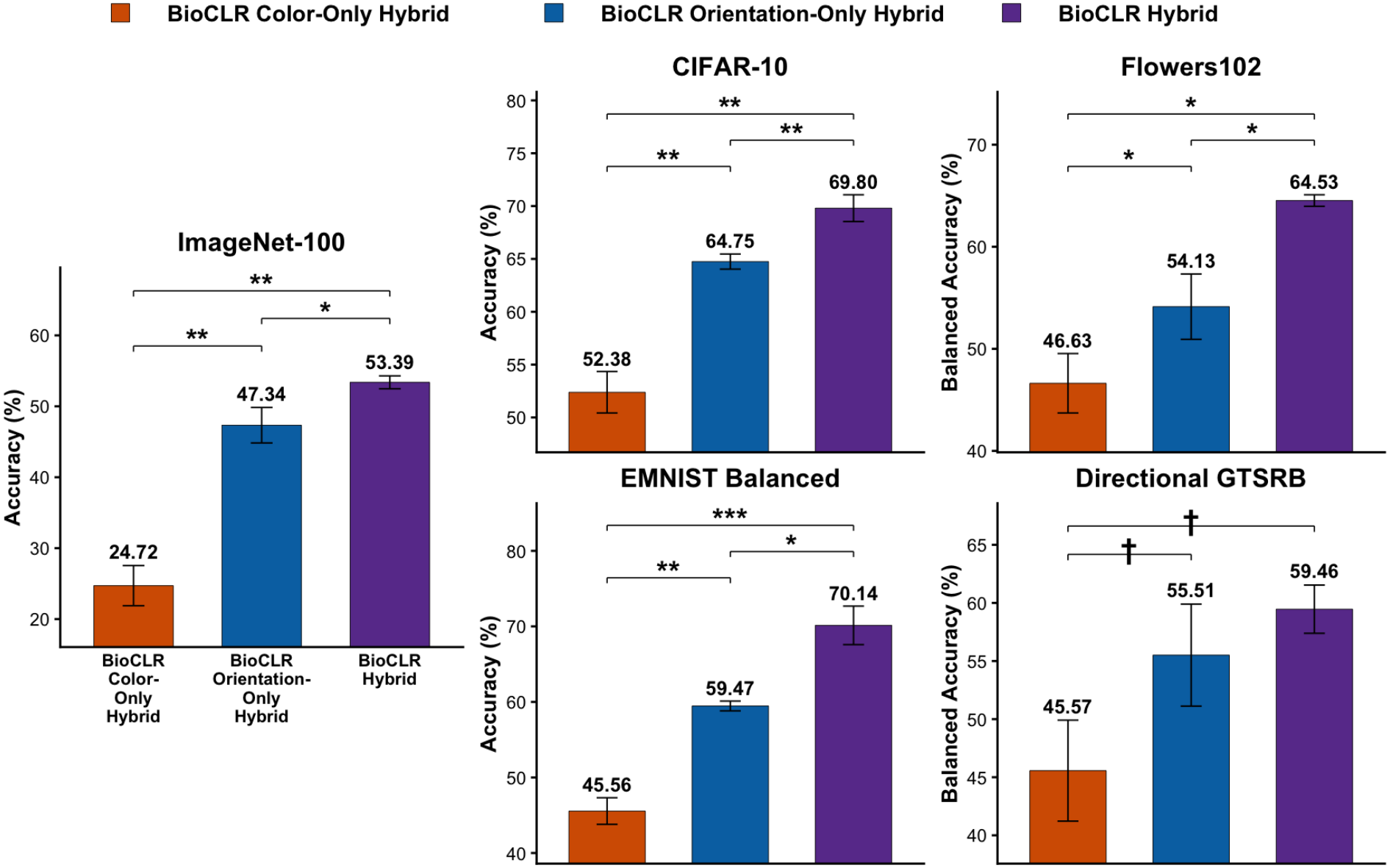
Frozen linear evaluation comparing the color-only, orientation-only, and full BioCLR hybrids. Bars show mean test accuracy or balanced accuracy across three independent runs and error bars show standard deviation. Statistical annotations are based on two-sided Welch’s *t*-tests with BH FDR correction across the three pairwise comparisons within each dataset. \**p*_BH_ *<* 0.05, \*\**p*_BH_ *<* 0.01, and \*\*\**p*_BH_ *<* 0.001; † indicates marginal significance (0.05 ≤ *p*_BH_ *<* 0.10).

Figure 7 and Table B.6 show the limited-label fine-tuning results. On ImageNet-100 with 1% labels (Figure 7, top panels), both the full and orientation-only models marginally outperform the color-only model, whereas the difference between the full and orientation-only models is not significant. With 10% labels, no significant differences are observed among the three models. On CIFAR-10, both the full and orientation-only models significantly outperform the color-only model at both label levels. The full model also achieves higher mean performance than the orientation-only model, with a marginal advantage at the 1% label level. On Flowers102, the full model significantly outperforms both single-teacher models at both the 1-shot and 5-shot levels, while the orientation-only model also significantly outperforms the color-only model. On EMNIST Balanced, no significant differences are observed among the three models with 0.1% labels. With 1% labels, both the full and orientation-only models significantly outperform the color-only model, whereas the difference between the full and orientation-only models is not significant. On Directional GTSRB, the orientation-only model achieves the highest mean performance at both the 5-shot and 50-shot levels, although differences among the three models are not statistically significant. With 50 shots per class, both the full and orientation-only models marginally outperform the color-only model.

**Figure 7:**
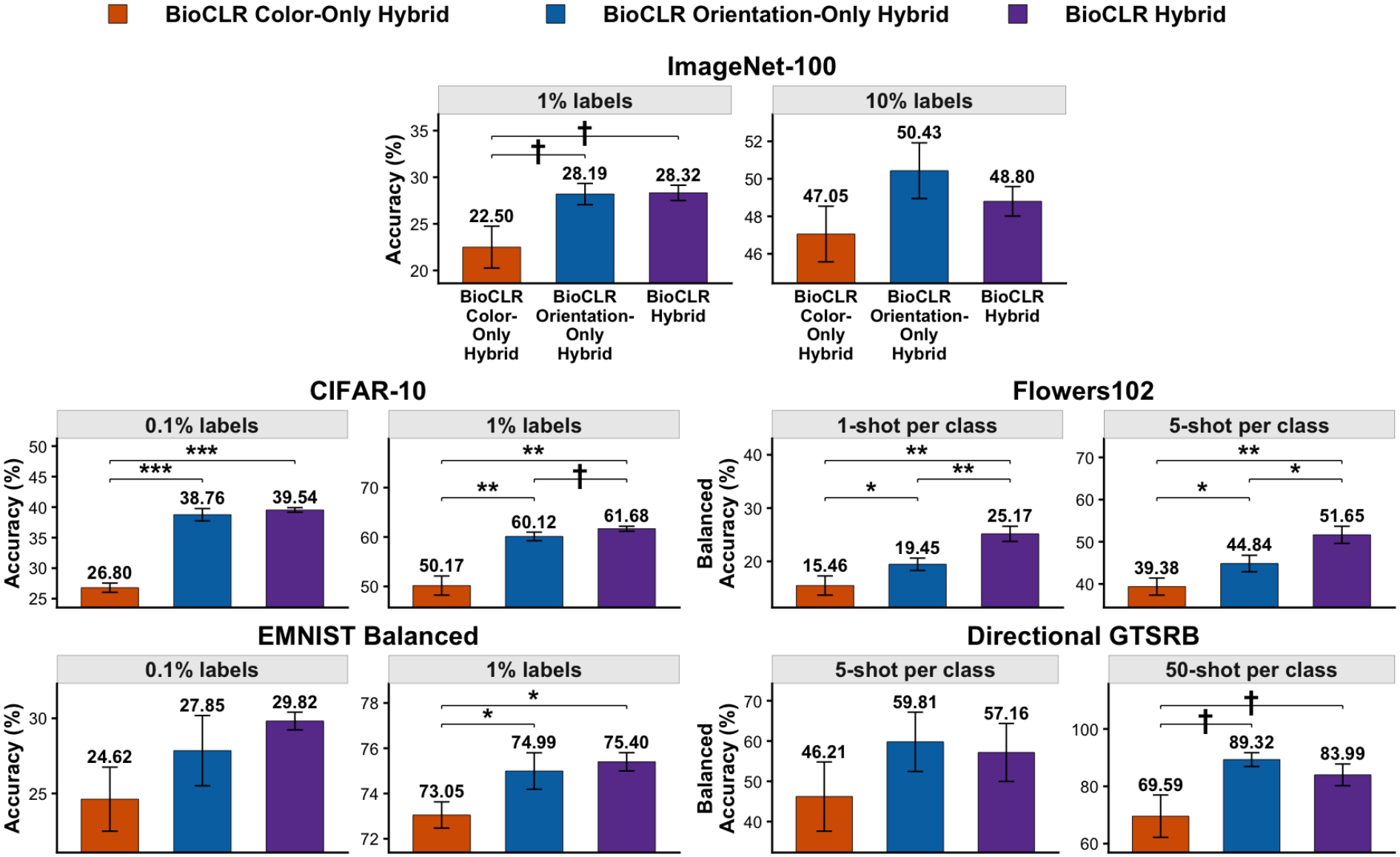
Limited-label fine-tuning comparing the color-only, orientation-only, and full BioCLR hybrids. Bars show mean test accuracy or balanced accuracy across three independent runs and error bars show standard deviation. Statistical annotations are based on two-sided Welch’s *t*-tests with BH FDR correction across the three pairwise comparisons within each dataset and label condition. \**p*_BH_ *<* 0.05, \*\**p*_BH_ *<* 0.01, and \*\*\**p*_BH_ *<* 0.001; † indicates marginal significance (0.05 ≤ *p*_BH_ *<* 0.10).

## 4 Discussion

In frozen linear evaluation, the BioCLR hybrid significantly outperformed standard SimCLR on ImageNet-100 and all four transfer datasets. This suggests that adding specialized teacher guidance to augmentation-based contrastive learning produces representations that are better for object recognition and that these benefits generalize beyond the pretraining dataset. However, it remains unclear whether these improvements result from using specialized color and orientation representations to guide learning or simply from providing additional color and orientation features to the learned visual representation. The post-hoc concatenation control addresses this concern. The BioCLR hybrid significantly outperformed this control across all five datasets. These results suggest that simply concatenating specialized representations after learning was insufficient to achieve the performance obtained through SSL guided by specialized teacher streams.

In limited-label fine-tuning, both standard SimCLR and the BioCLR hybrid generally outperformed supervised learning from scratch, suggesting that self-supervised pretraining improves the model’s adaptability to different object recognition tasks with limited labeled data (Chen et al., 2020b; Hénaff, 2020). Furthermore, the BioCLR hybrid significantly outperformed standard SimCLR in five transfer conditions, suggesting that specialized teacher guidance provides additional benefits for task adaptation. However, on ImageNet-100, the BioCLR hybrid showed only a marginal advantage over SimCLR with 1% labels and a marginally lower mean with 10% labels. These findings suggest that the benefits of specialized teacher guidance for limited-label fine-tuning are more apparent on transfer datasets than on the pretraining dataset.

The results on EMNIST Balanced suggest that specialized teacher guidance can mitigate the negative transfer observed with SimCLR. SimCLR performed significantly worse than the untrained baseline in frozen linear evaluation and than supervised learning from scratch in limited-label fine-tuning with 1% labels. This is consistent with previous findings that invariances learned through augmentation-based contrastive learning can harm downstream tasks that depend on the information altered by these augmentations (Xiao et al., 2021; Lee et al., 2021). In contrast, the BioCLR hybrid significantly outperformed SimCLR in both cases and did not show significant negative transfer.

Although the BioCLR hybrid significantly outperformed both the AugSelf and RotNet hybrids on ImageNet-100, it did not consistently achieve the best frozen linear evaluation performance across the transfer datasets. However, its advantages were more consistent during limited-label fine-tuning, where it achieved the highest mean performance across ImageNet-100 and the transfer datasets in eight of the ten conditions. These findings suggest that the benefits of incorporating attribute-related information into SSL models depend on how this information is learned and how the pretrained model is subsequently evaluated.

The Flowers102 results illustrate the distinction between frozen linear evaluation using the full labeled training set and limited-label fine-tuning using only a small number of labeled examples. The AugSelf hybrid significantly outperformed the BioCLR hybrid in frozen linear evaluation, whereas the BioCLR hybrid significantly outperformed the AugSelf hybrid after fine-tuning with five labeled examples per class and marginally outperformed it with one labeled example per class. These findings suggest that frozen linear evaluation using the full labeled training set does not fully predict how well a model will adapt to new tasks when labeled data are limited, extending previous findings that frozen linear evaluation does not fully predict fine-tuning performance (He et al., 2022; Kumar et al., 2022).

The ablation study clarifies the contributions of the two teachers. In both frozen linear evaluation and limited-label fine-tuning, the orientation-only model generally performed better than the color-only model. This is consistent with the effectiveness of rotation prediction as a self-supervised learning signal (Gidaris et al., 2018). However, color guidance still contributed: in frozen linear evaluation, the full BioCLR hybrid model significantly or marginally outperformed the color-only and orientation-only BioCLR hybrid models in nine of the ten pairwise comparisons across the five datasets. Therefore, combining color and orientation guidance improved frozen linear evaluation performance. The benefits of combining the two teachers were less consistent during limited-label fine-tuning: Flowers102 was the only dataset on which the full model significantly outperformed both single-teacher models, possibly because both color and orientation information are important for distinguishing fine-grained flower categories. These findings suggest that the contribution of each type of teacher guidance depends on the task and the type of evaluation.

From a neuroscience perspective, our findings suggest that specialized visual streams may provide learning signals that guide general-purpose visual representations needed for object recognition, beyond those gained from different views of the same object. These learning signals may help the visual system develop representations that can later support learning unfamiliar object categories from only a few labeled examples. Previous studies have suggested that specialized visual streams in the brain may arise because specialization improves performance on supervised visual tasks (Rueckl et al., 1989; Jacobs et al., 1991; Han & Sereno, 2024b). Our results extend this idea by suggesting that specialized streams may also support self-supervised learning through interactions between streams. The complementary benefits of the two teachers suggest that signals from multiple specialized streams can jointly support visual learning, with their relative contributions depending on the task. In particular, learning to distinguish categories that differ in both color and orientation may benefit more from guidance involving both attributes. Our findings also lead to the prediction that disrupting signals from color- or orientation-biased visual areas during learning could impair the development of object representations in other visual areas and affect visual recognition even after normal function in the disrupted areas is restored.

From a deep learning perspective, our findings suggest the importance of incorporating feature-specific visual information into SSL. The BioCLR hybrid illustrates a strategy in which multiple specialized representations provide complementary guidance to a shared model. This approach could be extended beyond color and orientation to other visual attributes and may help produce better representations that remain adaptable across tasks with limited supervision. At the same time, comparisons of the BioCLR hybrid with the AugSelf and RotNet hybrid models show that incorporating color- or orientation-related objectives through different learning mechanisms does not produce equally useful representations or equally adaptable models. These findings suggest that the design of the learning mechanism is as important as the choice of visual information incorporated into the learning objective.

## 5 Limitations

The current study has several limitations that may suggest directions for future work. First, the representational changes underlying the observed accuracy differences remain unclear. Specialized teacher guidance may preserve attribute information that receives less emphasis under standard SimCLR, alter how this information is organized in relation to object identity, or cause both effects. Attribute decoding and analyses of representational geometry, before and after fine-tuning, could distinguish these possibilities and help explain why a model can show greater adaptability during limited-label fine-tuning without achieving the best frozen linear evaluation accuracy. Second, BioCLR captures the general idea that specialized representations can provide learning signals to another visual representation, but it does not attempt to reproduce the development or complexity of specialized visual processing in the brain. The teacher networks are trained before the student and remain frozen, so the model does not address whether specialized and general-purpose representations develop sequentially or together through reciprocal interactions. The teacher objectives are also simplified: the color teacher classifies only the hue of the average color in the central region of an image, and the orientation teacher captures only one aspect of orientation processing. Future models could use more biologically realistic teacher objectives and allow specialized and general-purpose representations to develop jointly. In addition, future work may test whether the computational principles found in this study generalize across different architectures, pretraining datasets, forms of specialized information, and visual tasks.

## 6 Conclusion

In summary, in visual representation learning without human-annotated labels, our findings show that specialized visual streams can provide important learning signals that improve the performance of augmentation-based contrastive learning. First, adding color and orientation information to SimCLR through teacher guidance improved representation quality. Second, this guidance improved adaptability to various object recognition tasks when labeled data were limited. Third, how this information is incorporated also matters: compared with the AugSelf and RotNet hybrids (which encode color and orientation in other ways), the BioCLR hybrid achieved better representation quality on the pretraining dataset and generally better adaptability across datasets. Fourth, orientation guidance generally contributed more than color guidance, and combining the two provided complementary benefits, particularly in terms of representation quality.

## Author Contributions

Z.H.: Conceptualization, Data curation, Formal analysis, Investigation, Methodology, Software, Validation, Visualization, Writing – original draft, Writing – review & editing.

Y.L.: Writing – review & editing.

E.M.E.: Writing – review & editing.

A.B.S.: Funding acquisition, Methodology, Resources, Supervision, Visualization, Writing – review & editing.

## Acknowledgments

This work was supported in part by funds from Purdue University to A.B.S. and by the United States Air Force Office of Scientific Research under award no. FA9550-23-F-0014 to E.M.E.

## A Detailed methods

### A.1 Datasets

#### A.1.1 SSL pretraining dataset

ImageNet-100 is used for SSL pretraining. Defined by Tian et al. (2020a), it is a 100-class subset of the ImageNet dataset of natural object images (Deng et al., 2009), containing 126,689 training images and 5,000 validation images. We use a publicly released copy of this subset^1^. The training split is used for teacher network training and SSL pretraining of the student network. The same training split also provides the labeled examples used for subsequent frozen linear evaluation and limited-label fine-tuning on ImageNet-100. The validation split is used as the held-out in-domain evaluation test set. Example ImageNet-100 images are shown in Figure A.1.

#### A.1.2 Transfer datasets

Four transfer datasets are used to evaluate how well the SSL-pretrained representations and models transfer to new visual recognition tasks. For each transfer dataset, the official training split is used for frozen linear evaluation and limited-label fine-tuning training. The official test split is used for final evaluations.

CIFAR-10 is used as a near-domain transfer dataset because it also contains natural object images (Krizhevsky & Hinton, 2009). It contains 50,000 training images and 10,000 test images from 10 object categories. Example CIFAR-10 images are shown in Figure A.2.

Flowers102 is used to evaluate transfer to fine-grained object recognition (Nilsback & Zisserman, 2008), where distinctions among categories depend on color. It contains 102 flower categories. The current study uses the 1,020-image official training split for frozen linear evaluation and limited-label fine-tuning training, and the 6,149-image official test split for final evaluation. Example Flowers102 images are shown in Figure A.3.

EMNIST Balanced is used to evaluate transfer from natural object image representations to handwritten-character recognition (Cohen et al., 2017), where distinctions among categories depend on orientation. It contains 47 character classes, with 112,800 training images and 18,800 test images. Because EMNIST images are stored transposed, each image is first transposed to its upright orientation. The official training split is used for frozen linear evaluation and limited-label fine-tuning training, and the official test split is used for final evaluation. Example EMNIST Balanced images are shown in Figure A.4.

A directional subset of the German Traffic Sign Recognition Benchmark (GTSRB) is used to evaluate transfer to an object-recognition task in which both orientation and color information are relevant (Stallkamp et al., 2011). Eight directional classes are retained: dangerous curve to the left, dangerous curve to the right, turn right ahead, turn left ahead, go straight or right, go straight or left, keep right, and keep left. The resulting dataset contains 3,180 training images and 1,440 test images. The filtered training split is used for frozen linear evaluation and limited-label fine-tuning training, and the filtered test split is used for final evaluation. Example Directional GTSRB images are shown in Figure A.5.

### A.2 Image preprocessing

The same resizing and center-cropping procedure is applied to all datasets. Images are resized while preserving their aspect ratio so that the shorter side is 256 pixels, and are then center-cropped to 224 × 224. Therefore, smaller images are upsampled before the center crop, whereas larger images are downsampled as necessary. All model-specific augmentations during SSL training are applied after this common resizing and center-cropping procedure. EMNIST Balanced images are converted from a single grayscale channel to three identical channels by copying the same grayscale information across all three channels, so that they have the same number of channels as the color images in the other datasets.

Before images are provided to the neural networks, each channel is normalized using the same standard ImageNet channel mean and standard deviation. The channel mean vector, ordered as red, green, and blue, is

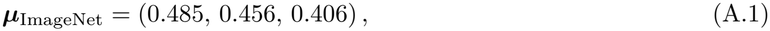

and the channel standard deviation vector is

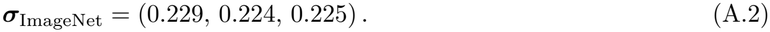

**Figure A_1:**
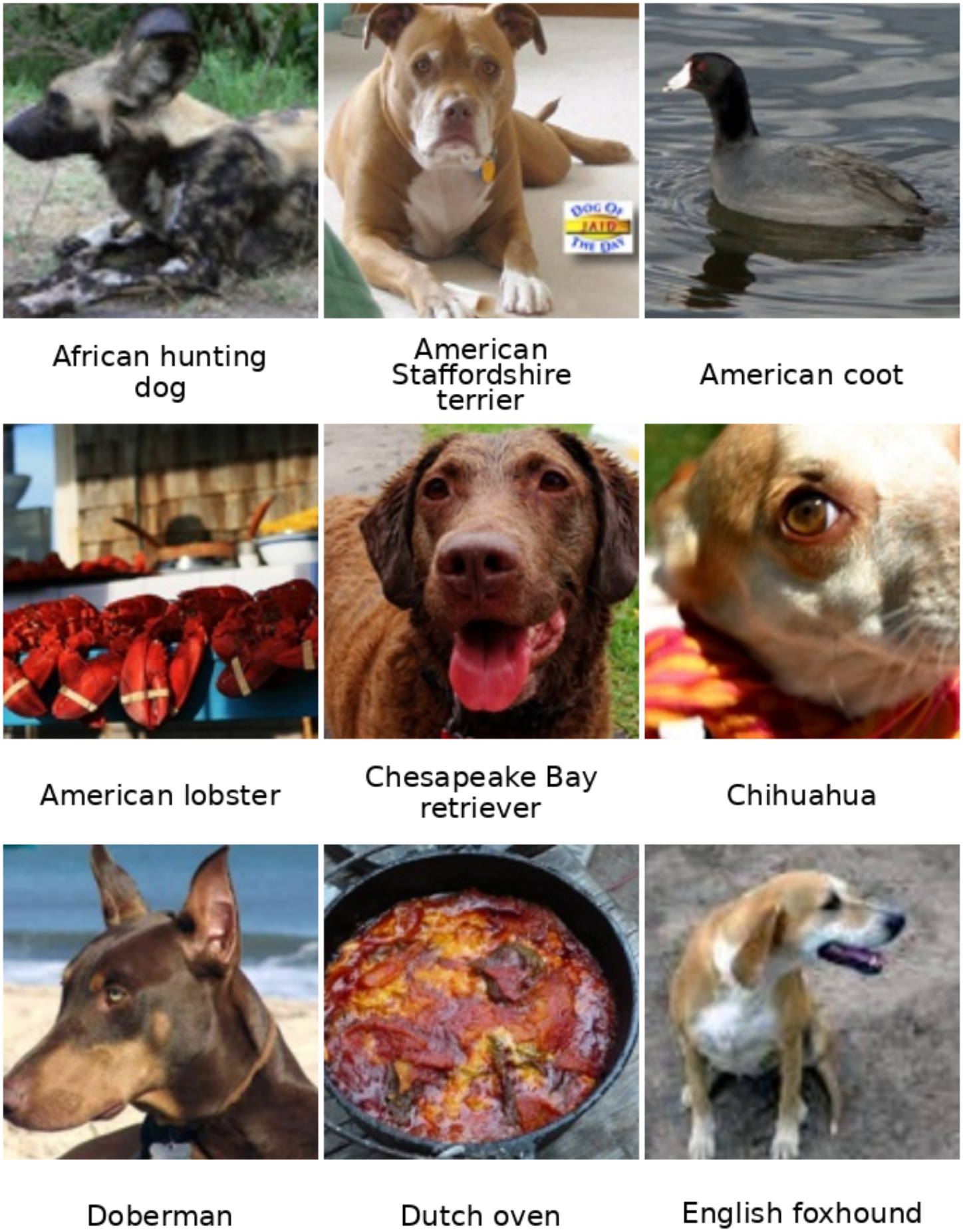
Example ImageNet-100 images after preprocessing.

Let *x*_*c,i,j*_ denote the pixel value at spatial location (*i, j*) in channel *c*, where *c* ∈ {*R, G, B*}, and let [***µ***_ImageNet_]_*c*_ and [***σ***_ImageNet_]_*c*_ denote the corresponding channel mean and standard deviation. Each pixel value is normalized as

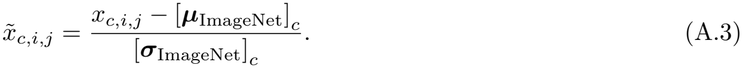

**Figure A_2:**
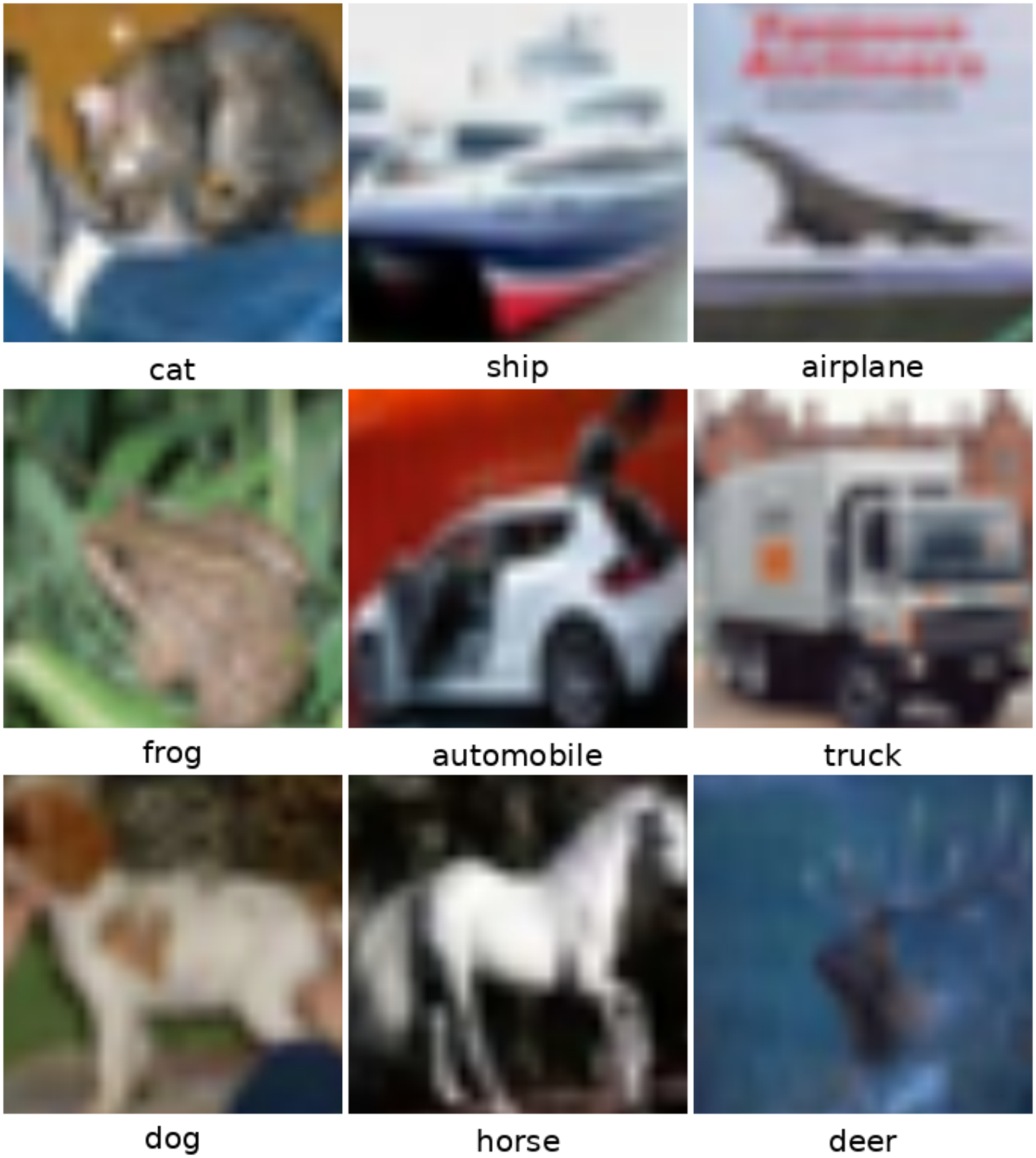
Example CIFAR-10 images after preprocessing.

### A.3 BioCLR hybrid model

#### A.3.1 BioCLR hybrid overview

We combine SimCLR with BioCLR so that the same student network is jointly shaped by both specialized teacher guidance and augmentation-based contrastive learning. We refer to this model as the SimCLR + BioCLR hybrid model (or BioCLR hybrid).

The BioCLR hybrid model trains the student encoder by jointly optimizing three loss functions: one SimCLR loss that encourages the student to develop representations that remain stable across transformations of the same image, and two BioCLR losses (one guided by the color teacher and the other by the orientation teacher) that encourage the student to retain color and orientation information emphasized by the specialized teacher encoders. The three losses are summed to form the BioCLR hybrid loss.

#### A.3.2 The SimCLR component

SimCLR learns visual representations by comparing the two independently augmented views generated from the same original preprocessed image. The two augmented views are processed by the student encoder and then mapped through a separate SimCLR projection head. The projected representations of the two views are used to calculate the SimCLR component loss. The SimCLR component is illustrated in Figure A.6(a). In the current study, each ImageNet-100 training image is preprocessed first and then each augmented view is independently generated from this preprocessed image using a random resized crop covering between 8% and 100% of its area, with an aspect ratio between 3*/*4 and 4*/*3. The crop is resized to 224 × 224. The following augmentations are then applied in order: horizontal flipping with a probability of 0.50; color jittering with a probability of 0.80; grayscale conversion with a probability of 0.20; and Gaussian blur with a probability of 0.50. During color jittering, four color adjustments are applied in random order: brightness, contrast, and saturation, with factors sampled uniformly from [0.2, 1.8], and hue, with a shift sampled uniformly from [−0.2, 0.2]. During Gaussian blur, each image is convolved with a 23 × 23 Gaussian kernel whose standard deviation is sampled uniformly from [0.1, 2.0]. This augmentation pipeline is applied independently twice to each preprocessed image to generate its two views.

**Figure A_3:**
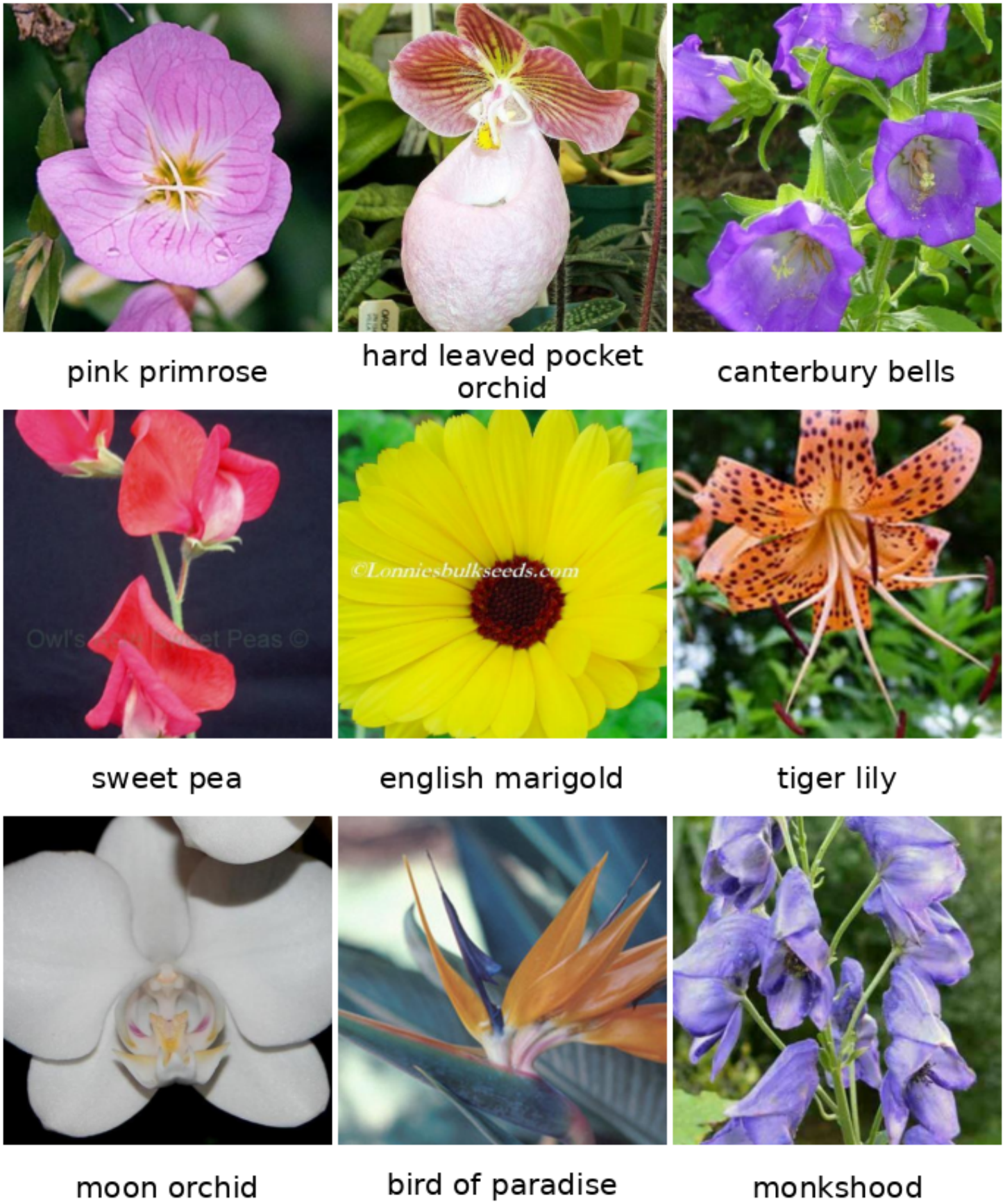
Example Flowers102 images after preprocessing.

**Figure A_4:**
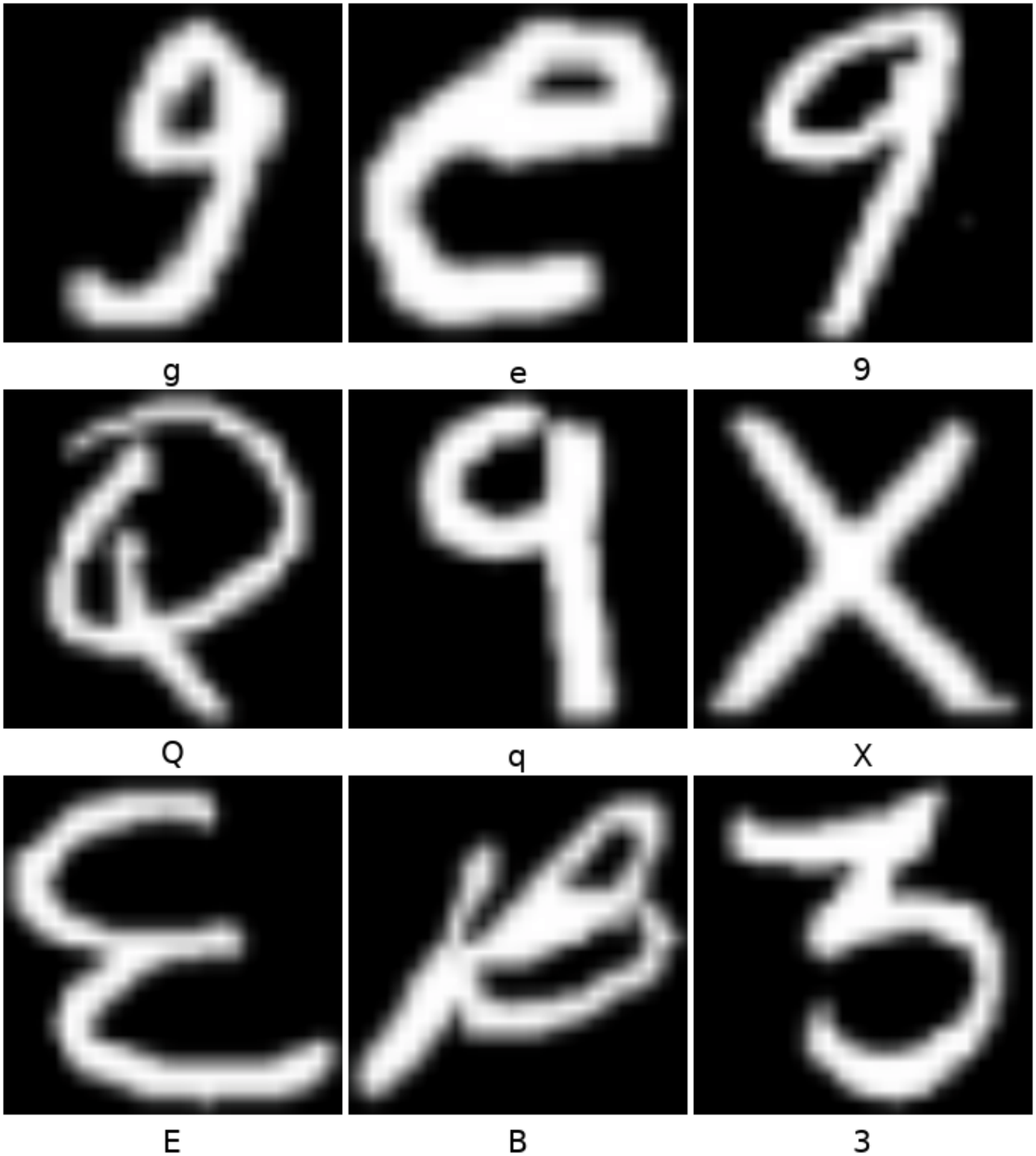
Example EMNIST Balanced images after preprocessing.

**Figure A_5:**
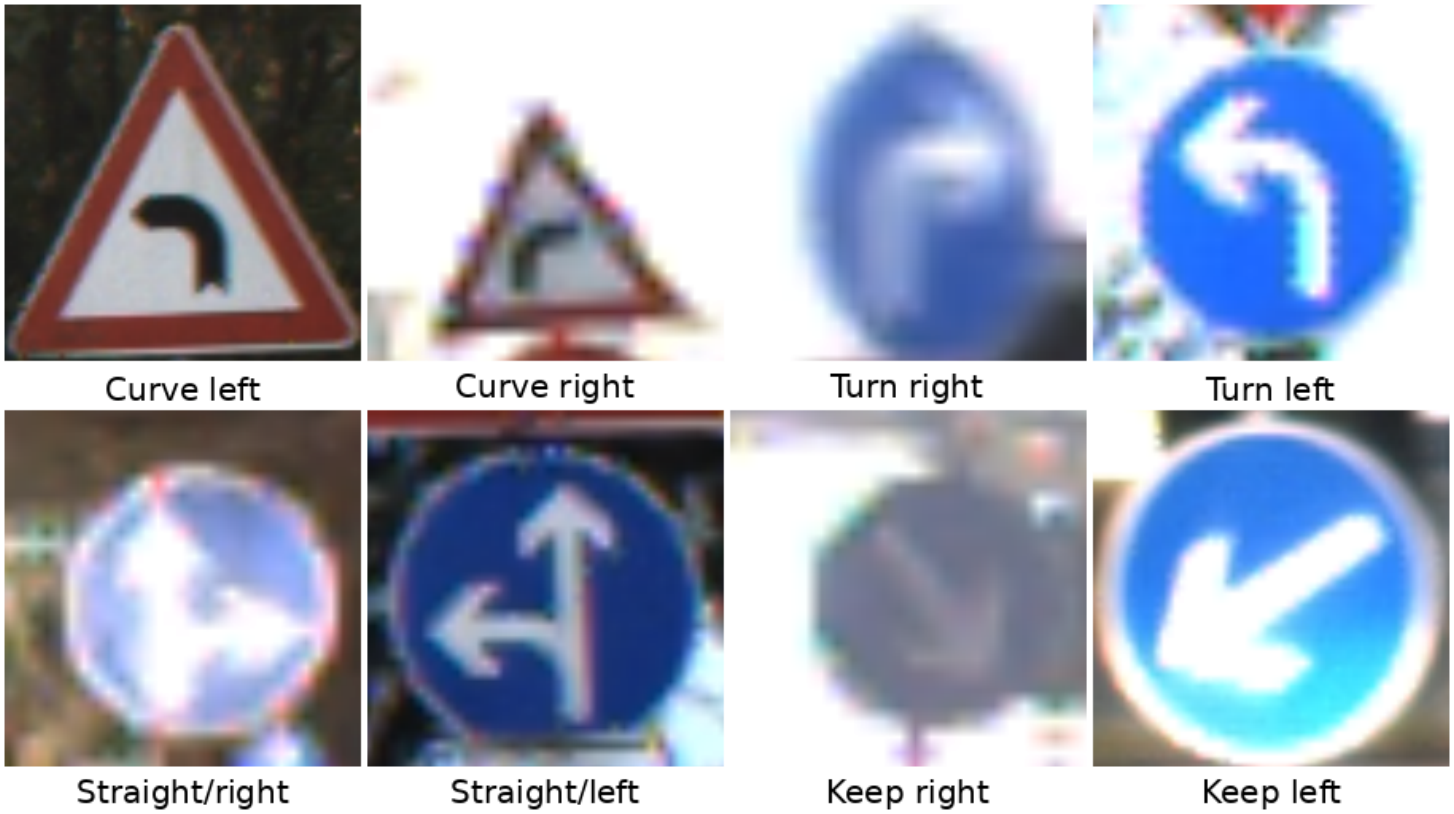
Example Directional GTSRB images after preprocessing.

**Figure A_6:**
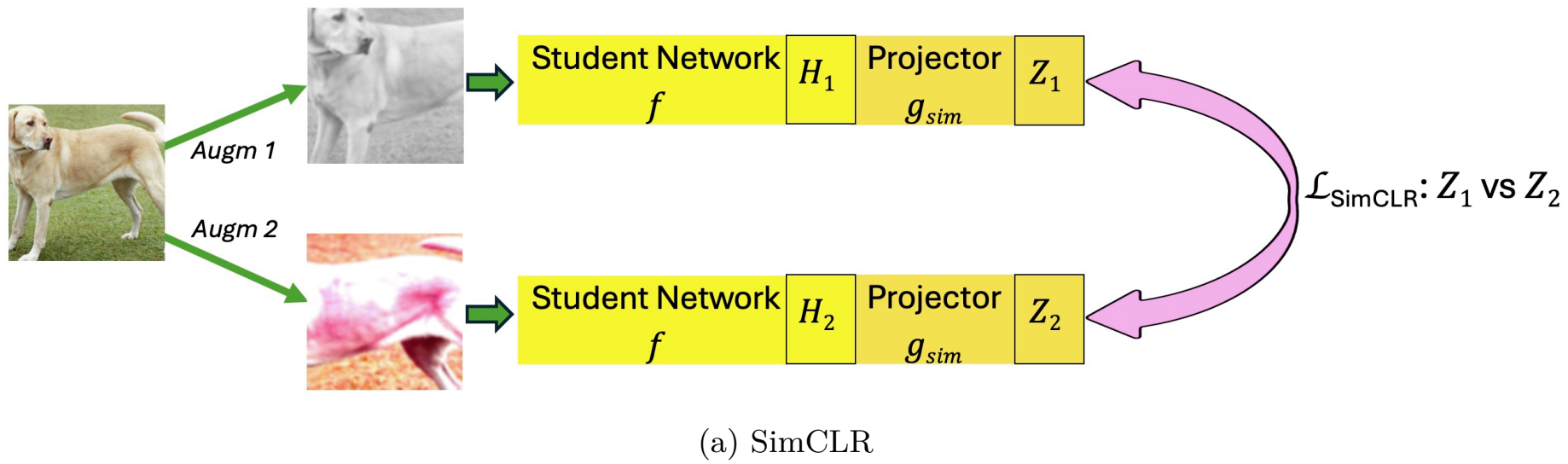
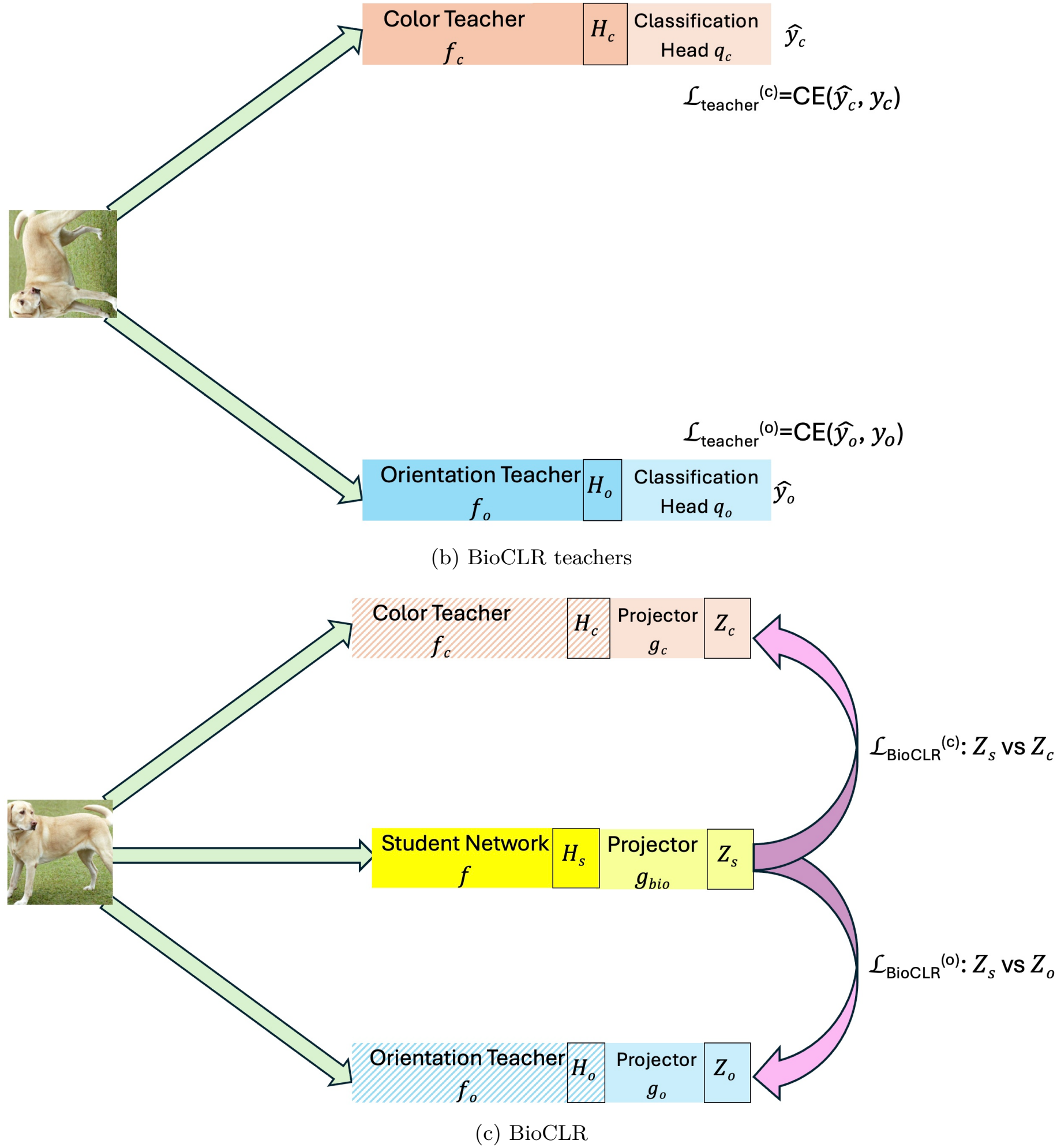
SimCLR, the BioCLR teachers and BioCLR. (a) SimCLR. Two augmented views of each image pass through the student network *f*. The SimCLR projector *g*_sim_ maps their representations *H*_1_ and *H*_2_ to the embeddings *Z*_1_ and *Z*_2_. The SimCLR loss (pink arrow; 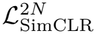 in standard SimCLR and the AugSelf and RotNet hybrids; *ℒ*_SimCLR_ in the BioCLR hybrid) compares *Z*_1_ and *Z*_2_. Both student networks share the weights *f*. (b) BioCLR teachers. Each randomly rotated image passes through the color teacher *f*_*c*_ and the orientation teacher *f*_*o*_. The classification heads *q*_*c*_ and *q*_*o*_ map their representations *H*_*c*_ and *H*_*o*_ to the predictions *ŷ*_*c*_ and *ŷ*_*o*_. Each teacher has its own cross-entropy (CE) loss: 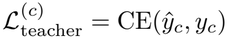 for the color teacher and 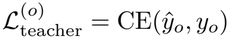 for the orientation teacher. (c) BioCLR. Each upright original preprocessed image passes through the student network *f* and the two teachers from (b). The teachers are now fixed (hatched). The projectors *g*_bio_, *g*_*c*_ and *g*_*o*_ map the representations *H*_*s*_, *H*_*c*_ and *H*_*o*_ to the embeddings *Z*_*s*_, *Z*_*c*_ and *Z*_*o*_. The BioCLR losses 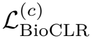 and 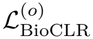 (pink arrows) compare *Z*_*s*_ with *Z*_*c*_ and with *Z*_*o*_. Dog images adapted from a photo by Elf and Djmirko (Wikimedia Commons, CC BY-SA 3.0).

#### A.3.3 The BioCLR component

The BioCLR component contains a color teacher network, an orientation teacher network, and a student network. The teacher networks are first trained to encode different visual attributes, and their encoders are then frozen during the training of the student network.

The BioCLR teacher training is illustrated in Figure A.6(b). During teacher training, each teacher network consists of an encoder and a classification head. The classification head maps the encoder features to a hidden representation, applies a ReLU activation, and then maps the hidden representation to the required number of classes for recognizing color or orientation. Only during the training of both teacher networks is each input image randomly rotated to one of four orientations (up, down, left, or right) each time it is retrieved.

The color teacher is trained to classify images based on the hue of the average color of their central region. A centered region spanning 60% of the image height and 60% of the image width is extracted, corresponding to approximately 36% of the total image area. The mean red, green, and blue values are calculated across this region and converted from RGB color space to HSV color space. The resulting hue value ranges from 0 to 1 and is divided into 12 equal intervals. Each image is assigned to the interval containing its hue value, producing one of 12 hue-category labels. The color teacher is trained using cross-entropy loss to classify each image into one of the 12 hue categories.

The orientation teacher is trained to determine the orientation of each image. The orientation teacher classifies each image into one of the four orientation classes and is trained using cross-entropy loss. This task encourages the encoder to preserve visual information useful for distinguishing image orientation.

After teacher training, the classification head of each teacher encoder is removed and only the trained teacher encoders are retained and frozen for subsequent self-supervised training of a student encoder. The BioCLR component is illustrated in Figure A.6(c).

#### A.3.4 BioCLR hybrid training

The BioCLR hybrid model is composed of the SimCLR and BioCLR components (Figure 1). During Bio-CLR hybrid training, images are processed in minibatches. The color and orientation teacher encoders are frozen throughout training, and only the student encoder and the four projection heads (the color-teacher, orientation-teacher, BioCLR, and SimCLR projection heads) are trained. For each minibatch, the SimCLR and BioCLR component losses are summed, and the student encoder and projection heads are updated jointly by minimizing this total loss.

For the SimCLR component, the student encoder processes two augmented views of each image, and a separate trainable SimCLR projection head maps the resulting encoder features into a contrastive embedding space. The SimCLR component loss encourages the projected representations of the two augmented views of the same image to be similar and those of different images to be dissimilar.

For the BioCLR component, the student encoder and both frozen teacher encoders receive the same upright original preprocessed images. Their encoder features are mapped into a contrastive embedding space using separate trainable projection heads. The student encoder uses a BioCLR projection head, while the color and orientation teacher encoders each use their own projection head. For each image in a minibatch, the BioCLR component loss encourages the projected student representation to be similar to the projected color- and orientation-teacher representations of the same image and dissimilar to the projected student representations of other images in the minibatch.

This training is self-supervised because the learning signals are derived from the images themselves through specialized teacher representations and augmented views, without requiring human-annotated labels. For additional details, the complete SSL training procedure for the BioCLR hybrid model (Figure 1) is summarized below in Algorithm A.1 (Section A.6). After training, all projection heads are removed, and only the trained student encoder is retained for subsequent evaluation.

### A.4 AugSelf hybrid model and RotNet hybrid model

To compare the BioCLR hybrid model with alternative approaches that actively encourage color and orientation encoding during SSL, we implement two additional hybrid models adapted from AugSelf (Lee et al., 2021) and RotNet (Gidaris et al., 2018). Both models combine the SimCLR objective with auxiliary objectives for color and orientation, using a shared encoder and separate task-specific heads. Their shared encoders have the same architecture and number of parameters as the student encoder in the BioCLR hybrid model, but neither model uses separately trained teacher encoders. Both alternative hybrid models use the same SimCLR augmentation pipeline described in Section A.3.2, with additional model-specific augmentations for their auxiliary learning objectives, as described below.

For the AugSelf hybrid model, two independently augmented views of the same original preprocessed image are generated, with each view also rotated, with a probability of 0.5, by one of four possible angles (0^*°*^, 90^*°*^, 180^*°*^, and 270^*°*^), following Lee et al. (2021). Both views are processed by the same shared encoder, producing one representation for each view. These representations are passed separately through the SimCLR projection head to compute the contrastive loss. The encoder representations of the two views are also concatenated and provided to two separate auxiliary predictors. One predictor estimates the differences in brightness, contrast, saturation, and hue-jitter parameters applied to the two views, using mean squared error loss. The other predictor classifies the orientation difference between the views into one of four categories, using cross-entropy loss. The SimCLR loss and both auxiliary losses jointly update the shared encoder.

For the RotNet hybrid model, two independently augmented views of the same original preprocessed image are processed by a shared encoder, and their representations are passed through a SimCLR projection head to compute the contrastive loss. In a separate auxiliary branch, the original preprocessed image is randomly rotated to one of four orientations (up, down, left, right) each time it is retrieved. The rotated image is processed by the same shared encoder, and its representation is passed through two separate auxiliary predictors: one predicts the image’s orientation, and the other predicts its color category. The color category is defined by dividing the hue of the mean RGB color in the central 60% × 60% region into 12 categories, following the same procedure used for the BioCLR color teacher. Both auxiliary tasks use cross-entropy loss, and their losses are combined with the SimCLR loss to jointly update the shared encoder.

After training, all projection heads and auxiliary predictors are removed, and only the trained shared encoder is retained for subsequent evaluation.

### A.5 BioCLR hybrid ablation models

For the two ablation models of the BioCLR hybrid, both models retain the same student encoder and SimCLR component as the full BioCLR hybrid model. The color-only hybrid uses only the color-guided BioCLR loss, whereas the orientation-only hybrid uses only the orientation-guided BioCLR loss. The remaining training procedures and settings are unchanged.

### A.6 Training algorithm summary for BioCLR, AugSelf, and RotNet hybrids

Algorithm A.1 summarizes the training algorithm for all three hybrid models, where *m* denotes the model condition. All three hybrid models jointly optimize a SimCLR contrastive loss and two additional losses related to color and orientation. The network architectures and losses they use are detailed below in Sections A.7 and A.8.

#### Networks

In all three hybrid models, *f* denotes the encoder (the student encoder in the BioCLR hybrid), and *g*_sim_ denotes the SimCLR projection head attached to the encoder. The BioCLR hybrid uses the fixed color- and orientation-teacher encoders *f*_*c*_ and *f*_*o*_, and three projection heads: *g*_bio_ for the student, *g*_*c*_ for the color teacher, and *g*_*o*_ for the orientation teacher. The AugSelf and RotNet hybrids use the auxiliary color and orientation predictors *h*_*c*_ and *h*_*o*_.

#### Data and representations

*X* denotes a minibatch of *N* original preprocessed images, and 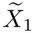 and 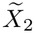 denote its two augmented views. *H*_1_ and *H*_2_ denote the encoder representations of the two views, and *Z*_1_ and *Z*_2_ denote their projected embeddings, whose *i*-th rows are **z**_1,*i*_ and **z**_2,*i*_. The remaining variables are model-specific:

- BioCLR hybrid: *H*_*s*_, *H*_*c*_, and *H*_*o*_ denote the student, color-teacher, and orientation-teacher encoder representations of *X*, and *Z*_*s*_, *Z*_*c*_, and *Z*_*o*_ denote their projected embeddings, whose *i*-th rows are **z**_*s,i*_ and **z**_*t,i*_, where *t* ∈ {*c, o*} denotes the color or orientation teacher.
- AugSelf hybrid: [·, ·] denotes concatenation along the feature dimension, and *P* denotes the 2*N* concatenated representation pairs. For each pair, Δ**c** denotes the differences in normalized color-jitter parameters, and Δ*o* denotes the orientation difference (modulo 4) between the first and second views.
- RotNet hybrid: *X*_*o*_ denotes a randomly rotated copy of *X, y*_*c*_ and *y*_*o*_ denote its color and orientation labels, and *H*_aux_ denotes its encoder representations.

#### Losses

*ℒ*_sim_, *ℒ*_*c*_, and *ℒ*_*o*_ denote the SimCLR, color, and orientation losses of each hybrid model, respectively, and *ℒ* denotes their sum. The BioCLR hybrid uses a modified NT-Xent loss, *ℒ*_SimCLR_ (Section A.8.1), whereas the AugSelf and RotNet hybrids use the standard NT-Xent loss, 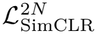 (Section A.8.2). 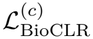 and 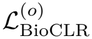 denote the BioCLR loss with the color and orientation teachers, respectively (Section A.8.1). MSE and CE denote the mean squared error loss and the cross-entropy loss.

### A.7 Architectures

#### A.7.1 Encoder architecture

All BioCLR teacher and student encoders use the standard ImageNet-scale ResNet-18 architecture (He et al., 2016). The final fully connected classification layer is replaced with an identity mapping, so that each encoder produces a 512-dimensional representation. The AugSelf hybrid and RotNet hybrid models also use the same encoder architecture.

#### A.7.2 Projection-head and auxiliary-predictor architectures

All three hybrid models (BioCLR, AugSelf, and RotNet) use the same architecture for their SimCLR projection heads. Each projection head consists of a two-layer multilayer perceptron that maps the 512-dimensional encoder representation to a 512-dimensional hidden representation, followed by a ReLU activation and a second fully connected layer that produces a 128-dimensional projected embedding. The projection-head parameters are trained separately for each model.

##### Algorithm A.1

SSL training algorithm for the BioCLR, AugSelf, and RotNet hybrid models. All notation is defined in Section A.6.

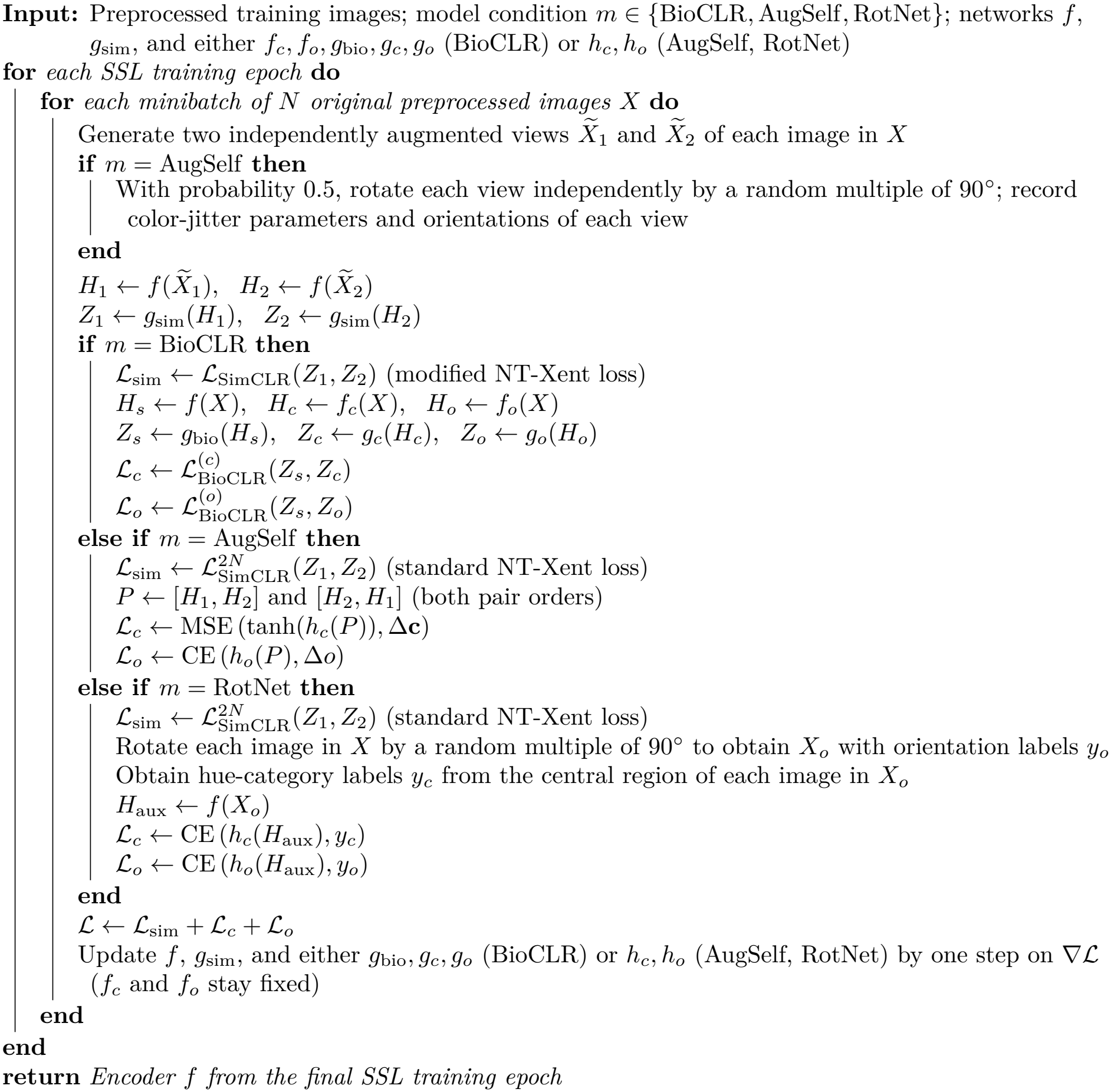

The BioCLR hybrid model additionally uses three projection heads with the same architecture: one for the student representation, one for the color teacher representation, and one for the orientation teacher representation. All four projection heads in the BioCLR hybrid model have separate parameters.

The AugSelf and RotNet hybrid models each additionally use two auxiliary predictors, a color predictor and an orientation predictor. In the AugSelf hybrid model, following the design of the original AugSelf model (Lee et al., 2021), each predictor receives the 1024-dimensional concatenated encoder representation of the two augmented views and passes it through two hidden layers with 512 units (each followed by batch normalization and a ReLU activation) and an output layer with four units (four color-jitter parameters or four orientation classes). In the RotNet hybrid model, each predictor has the same architecture as the classification heads of the BioCLR teachers: it receives the 512-dimensional encoder representation of the rotated image and passes it through one hidden layer with 512 units and a ReLU activation, followed by an output layer with 12 units for the color predictor and four units for the orientation predictor.

### A.8 Loss functions

#### A.8.1 BioCLR hybrid model loss

The BioCLR hybrid loss is the summation of a SimCLR loss and two BioCLR losses (one guided by the color teacher and the other guided by the orientation teacher).

The SimCLR loss in the BioCLR hybrid is modified from the standard NT-Xent loss so that negative pairs are formed only with representations from the opposite augmented-view batch. This ensures that both the BioCLR and the SimCLR losses in the BioCLR hybrid model use the same number of negative pairs (*N* − 1) per anchor image. Let **z**_1,*i*_ and **z**_2,*i*_ denote the projected representations of the two augmented views of image *i*. For each representation **z**_1,*i*_ in the first batch, the corresponding representation **z**_2,*i*_ in the second batch is the positive, whereas representations **z**_2,*j*_ from other images (*j* ≠ *i*) are negatives. The loss is computed in both directions and averaged:

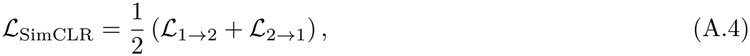

where the loss from the first batch to the second batch is defined as

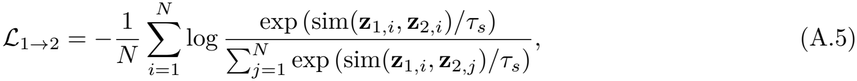

where *τ*_*s*_ is the SimCLR temperature parameter. The loss in the other direction, *ℒ*_2*→*1_, is defined analogously by exchanging the two augmented-view batches.

For the BioCLR losses, each projected student representation is compared with the corresponding projected teacher representation as a positive pair and the projected student representations of other images in the minibatch as negative pairs. For each image *i* in a minibatch of *N* images, let **z**_*s,i*_ denote its projected student representation and **z**_*t,i*_ denote its projected representation from teacher *t*. The BioCLR loss for each teacher is defined as

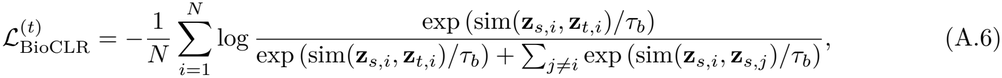

where *τ*_*b*_ is the BioCLR temperature parameter and sim(·, ·) denotes the cosine similarity between two vectors.

The BioCLR hybrid model combines the SimCLR loss with the color- and orientation-guided BioCLR losses:

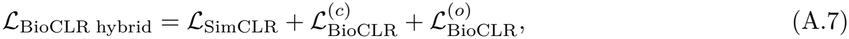

where 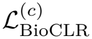 and 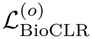 denote the BioCLR loss computed with the color teacher (*t* = *c*) and the orientation teacher (*t* = *o*), respectively.

#### A.8.2 AugSelf hybrid model and RotNet hybrid model losses

The SimCLR components of the AugSelf and RotNet hybrid models use the standard NT-Xent loss, denoted 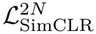, in which the two augmented-view batches are combined and each representation is contrasted with all other representations in the combined batch except itself.

The AugSelf hybrid model combines the SimCLR loss with two auxiliary losses. The color loss uses mean squared error to predict differences in brightness, contrast, saturation, and hue augmentation parameters between two augmented views. The orientation loss uses cross-entropy to classify the orientation difference between the two views into one of four categories. The total loss is

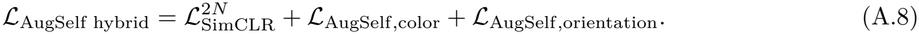

The RotNet hybrid model combines the SimCLR loss with two auxiliary classification losses. Both are cross-entropy losses on the rotated image: the color loss for classifying it into one of 12 hue categories based on the average hue in its central region, and the orientation loss for classifying it into one of four orientations. The total loss is

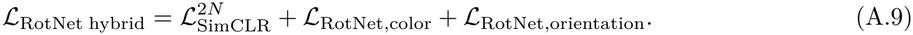

#### A.8.3 BioCLR hybrid ablation model loss

In the ablation study, the color-only and orientation-only BioCLR hybrid models are trained with

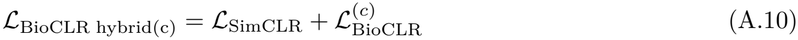

and

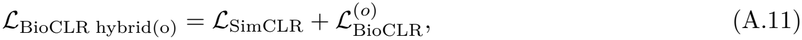

respectively.

### A.9 Model evaluation

After SSL training, the encoder of each model is evaluated on each dataset using frozen linear evaluation (Section A.9.1) and limited-label fine-tuning (Section A.9.2). No data augmentation is applied during model evaluation. The entire experiment is repeated in three independent runs. In each run, all networks are trained from new random initializations without fixed random seeds, and all pretraining and evaluation steps are repeated independently.

#### A.9.1 Frozen linear evaluation

During frozen linear evaluation, the encoder is fixed, and only the newly initialized linear classification head is trained using cross-entropy loss. This procedure is applied to all SSL-pretrained models and an untrained baseline, whose randomly initialized encoder is also kept fixed. For the post-hoc concatenation control, the 512-dimensional representations from independently trained standard SimCLR, color-teacher, and orientation-teacher encoders are concatenated into a 1536-dimensional representation. All three encoders are kept fixed, and only a linear classification head is trained on the concatenated representations. The complete labeled training split of each dataset is used. After training, the models are evaluated on the held-out test split. The ImageNet-100 validation split is used for ImageNet-100, whereas the official test splits are used for CIFAR-10, Flowers102, EMNIST Balanced, and Directional GTSRB.

#### A.9.2 Limited-label fine-tuning

During limited-label fine-tuning, a newly initialized linear classification head is attached to the encoder, and both the encoder and classification head are updated using cross-entropy loss. For ImageNet-100, fine-tuning uses 1% and 10% of the available training images. CIFAR-10 uses 0.1% (5 images per class) and 1% (50 images per class) of the training images, and EMNIST Balanced uses 0.1% (2 images per class) and 1% (24 images per class) of the training images. Flowers102 uses one or five labeled training images per class, whereas Directional GTSRB uses 5 or 50 labeled images per class. Labeled subsets are sampled randomly within each class. Within each run, all models are fine-tuned on the same subset, and a new subset is sampled in each run. After fine-tuning, the encoder and classification head from the final training epoch are evaluated on the same held-out test splits used for frozen linear evaluation.

The supervised baseline uses the same encoder architecture with a linear classification head attached. It is trained from scratch on the same labeled subset with the same fine-tuning loss, number of epochs, and batch size. Because it has no pretrained weights, it uses a larger encoder learning rate and updates its BatchNorm statistics during training (Section A.10).

#### A.9.3 Evaluation metrics

Classification accuracy (the percentage of correctly classified test images) is reported for ImageNet-100, CIFAR-10, and EMNIST Balanced. To account for differences in class frequencies, balanced accuracy (the mean of the per-class accuracies) is reported for Flowers102 and Directional GTSRB.

### A.10 Optimization and training settings

All teacher training, SSL training, frozen linear evaluation, and limited-label fine-tuning use a fixed number of training epochs. The model from the final epoch is retained. All models are optimized using Adam with a cosine annealing learning-rate schedule and a weight-decay coefficient of 1 × 10^*−*4^.

The color and orientation teacher networks are trained for 200 epochs with a batch size of 256 and an initial learning rate of 1 × 10^*−*3^. SSL models are trained for 400 epochs with a batch size of 1024 and an initial learning rate of 1×10^*−*3^. The SimCLR temperature is *τ*_*s*_ = 0.50 in the standard SimCLR and in the SimCLR components of all hybrid models. The BioCLR temperature is *τ*_*b*_ = 0.10. During limited-label fine-tuning of SSL-trained encoders, BatchNorm layers are kept in evaluation mode.

Frozen linear evaluation classifiers are trained for 200 epochs with a batch size of 256 and an initial learning rate of 1 × 10^*−*2^. Limited-label fine-tuning is also performed for 200 epochs with a batch size of 256. For SSL-trained encoders, the initial learning rate is 1 × 10^*−*5^ for the encoder and 1 × 10^*−*3^ for the classification head. For the supervised baseline trained from random initialization, both the encoder and classification head use an initial learning rate of 1 × 10^*−*3^.

## B Result tables

In all tables, values are mean ± SD test accuracy (ImageNet-100, CIFAR-10, and EMNIST Balanced) or balanced accuracy (Flowers102 and Directional GTSRB) across three independent runs (%). Boldface indicates the highest mean in each row. Significant differences are summarized using statistical groupings (compact letter display; Piepho, 2004) based on two-sided Welch’s *t*-tests with BH FDR correction across all pairwise comparisons within each row (six comparisons in Table B.1 and three in the other tables; *p*_BH_ *<* 0.05). Each value is assigned one or more superscript letters (*a, b, c*, etc.), such that two values sharing at least one letter are not significantly different, whereas two values with no letter in common are significantly different. A value with more than one letter belongs to more than one group. For example, a value labeled *ab* does not differ significantly from values labeled *a* or from values labeled *b*, even though values labeled *a* and *b* differ significantly from each other. Letters are assigned separately for each row and cannot be compared across rows. Marginal differences are not shown in the tables.

**Table B.1:** Frozen linear evaluation comparing an untrained baseline, standard SimCLR, a post-hoc concatenation control, and the BioCLR hybrid. Values and statistical groupings are described at the beginning of Appendix B.

| Dataset | Untrained<br>Baseline | SimCLR | Post-hoc<br>Concatenation Control | BioCLR Hybrid |
|---|---|---|---|---|
| ImageNet-100 | 16.45 ± 0.69<sup>d</sup> | 49.29 ± 0.19<sup>c</sup> | 50.12 ± 0.20<sup>b</sup> | **53.39 ± 0.91<sup>a</sup>** |
| CIFAR-10 | 47.54 ± 0.65<sup>d</sup> | 65.14 ± 0.25<sup>c</sup> | 66.68 ± 0.29<sup>b</sup> | **69.80 ± 1.27<sup>a</sup>** |
| Flowers102 | 24.81 ± 1.38<sup>d</sup> | 58.91 ± 0.14<sup>b</sup> | 53.10 ± 0.23<sup>c</sup> | **64.53 ± 0.56<sup>a</sup>** |
| EMNIST Balanced | **72.65 ± 1.96<sup>a</sup>** | 40.55 ± 1.02<sup>c</sup> | 52.31 ± 0.63<sup>b</sup> | 70.14 ± 2.55<sup>a</sup> |
| Directional GTSRB | 47.60 ± 7.50<sup>abc</sup> | 35.94 ± 0.98<sup>c</sup> | 50.94 ± 1.70<sup>b</sup> | **59.46 ± 2.07<sup>a</sup>** |

**Table B.2:** Limited-label fine-tuning comparing a supervised baseline, standard SimCLR, and the BioCLR hybrid. Values and statistical groupings are described at the beginning of Appendix B.

| Dataset | Labels | Supervised Baseline | SimCLR | BioCLR Hybrid |
|---|---|---|---|---|
| ImageNet-100 | 1% labels | 11.23 ± 0.18<sup>*b*</sup> | 25.96 ± 1.49<sup>*a*</sup> | **28.32 ± 0.82<sup>*a*</sup>** |
|  | 10% labels | 31.39 ± 0.56<sup>*b*</sup> | **50.37 ± 0.21<sup>*a*</sup>** | 48.80 ± 0.79<sup>*a*</sup> |
| CIFAR-10 | 0.1% labels | 23.14 ± 0.83<sup>*b*</sup> | 35.52 ± 3.14<sup>*a*</sup> | **39.54 ± 0.38<sup>*a*</sup>** |
|  | 1% labels | 38.68 ± 1.70<sup>*c*</sup> | 57.92 ± 1.44<sup>*b*</sup> | **61.68 ± 0.50<sup>*a*</sup>** |
| Flowers102 | 1-shot per class | 14.90 ± 0.73<sup>*c*</sup> | 18.25 ± 0.47<sup>*b*</sup> | **25.17 ± 1.42<sup>*a*</sup>** |
|  | 5-shot per class | 35.15 ± 0.84<sup>*c*</sup> | 43.42 ± 1.82<sup>*b*</sup> | **51.65 ± 2.03<sup>*a*</sup>** |
| EMNIST Balanced | 0.1% labels | 20.74 ± 0.75<sup>*c*</sup> | 26.07 ± 1.33<sup>*b*</sup> | **29.82 ± 0.59<sup>*a*</sup>** |
|  | 1% labels | **76.93 ± 0.82<sup>*a*</sup>** | 72.26 ± 0.39<sup>*b*</sup> | 75.40 ± 0.40<sup>*a*</sup> |
| Directional GTSRB | 5-shot per class | 29.51 ± 6.58<sup>*b*</sup> | 55.80 ± 7.06<sup>*a*</sup> | **57.16 ± 7.20<sup>*a*</sup>** |
|  | 50-shot per class | 54.78 ± 5.88<sup>*b*</sup> | 82.48 ± 6.78<sup>*a*</sup> | **83.99 ± 3.79<sup>*a*</sup>** |

**Table B.3:**
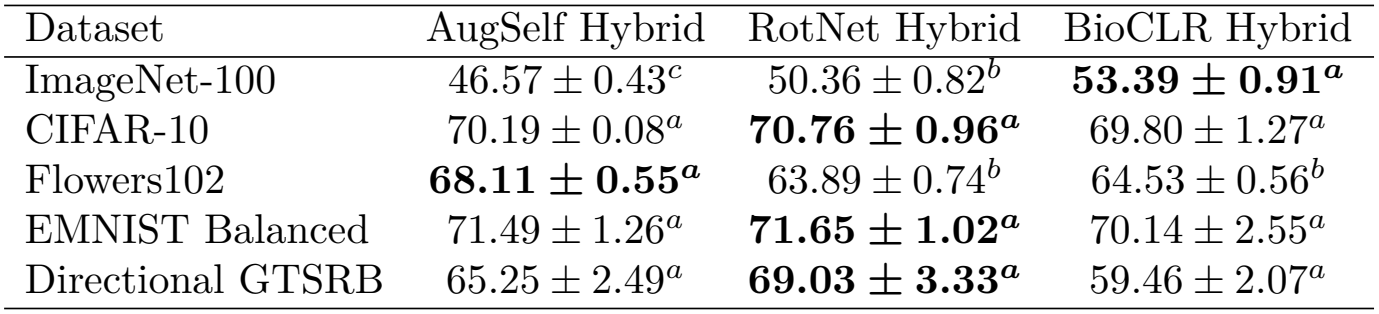
Frozen linear evaluation comparing the AugSelf, RotNet, and BioCLR hybrids. Values and statistical groupings are described at the beginning of Appendix B.

**Table B.4:** Limited-label fine-tuning comparing the AugSelf, RotNet, and BioCLR hybrids. Values and statistical groupings are described at the beginning of Appendix B.

| Dataset | Labels | AugSelf Hybrid | RotNet Hybrid | BioCLR Hybrid |
|---|---|---|---|---|
| ImageNet-100 | 1% labels | 21.92 ± 0.47<sup>*c*</sup> | 24.85 ± 0.98<sup>*b*</sup> | **28.32 ± 0.82<sup>*a*</sup>** |
|  | 10% labels | 43.89 ± 0.11<sup>*b*</sup> | 44.35 ± 1.17<sup>*b*</sup> | **48.80 ± 0.79<sup>*a*</sup>** |
| CIFAR-10 | 0.1% labels | 37.04 ± 2.25<sup>*a*</sup> | 39.14 ± 2.12<sup>*a*</sup> | **39.54 ± 0.38<sup>*a*</sup>** |
|  | 1% labels | 56.71 ± 1.19<sup>*b*</sup> | 58.10 ± 1.34<sup>*b*</sup> | **61.68 ± 0.50<sup>*a*</sup>** |
| Flowers102 | 1-shot per class | 19.11 ± 2.04<sup>*a*</sup> | 23.67 ± 2.45<sup>*a*</sup> | **25.17 ± 1.42<sup>*a*</sup>** |
|  | 5-shot per class | 42.69 ± 1.70<sup>*b*</sup> | 46.11 ± 2.65<sup>*ab*</sup> | **51.65 ± 2.03<sup>*a*</sup>** |
| EMNIST Balanced | 0.1% labels | **33.55 ± 2.30<sup>*a*</sup>** | 31.22 ± 2.15<sup>*a*</sup> | 29.82 ± 0.59<sup>*a*</sup> |
|  | 1% labels | 72.72 ± 0.82<sup>*b*</sup> | 73.37 ± 0.43<sup>*b*</sup> | **75.40 ± 0.40<sup>*a*</sup>** |
| Directional GTSRB | 5-shot per class | 49.96 ± 5.41<sup>*a*</sup> | **58.46 ± 3.77<sup>*a*</sup>** | 57.16 ± 7.20<sup>*a*</sup> |
|  | 50-shot per class | 80.09 ± 3.20<sup>*a*</sup> | 82.79 ± 6.70<sup>*a*</sup> | **83.99 ± 3.79<sup>*a*</sup>** |

**Table B.5:** Frozen linear evaluation comparing the color-only, orientation-only, and full BioCLR hybrids. Values and statistical groupings are described at the beginning of Appendix B.

| Dataset | BioCLR<br>Color-Only Hybrid | BioCLR<br>Orientation-Only Hybrid | BioCLR Hybrid |
|---|---|---|---|
| ImageNet-100 | 24.72 ± 2.83<sup>*c*</sup> | 47.34 ± 2.50<sup>*b*</sup> | **53.39 ± 0.91<sup>*a*</sup>** |
| CIFAR-10 | 52.38 ± 1.96<sup>*c*</sup> | 64.75 ± 0.71<sup>*b*</sup> | **69.80 ± 1.27<sup>*a*</sup>** |
| Flowers102 | 46.63 ± 2.91<sup>*c*</sup> | 54.13 ± 3.20<sup>*b*</sup> | **64.53 ± 0.56<sup>*a*</sup>** |
| EMNIST Balanced | 45.56 ± 1.76<sup>*c*</sup> | 59.47 ± 0.65<sup>*b*</sup> | **70.14 ± 2.55<sup>*a*</sup>** |
| Directional GTSRB | 45.57 ± 4.34<sup>*a*</sup> | 55.51 ± 4.39<sup>*a*</sup> | **59.46 ± 2.07<sup>*a*</sup>** |

**Table B.6:** Limited-label fine-tuning comparing the color-only, orientation-only, and full BioCLR hybrids. Values and statistical groupings are described at the beginning of Appendix B.

| Dataset | Labels | BioCLR<br>Color-Only Hybrid | BioCLR<br>Orientation-Only Hybrid | BioCLR Hybrid |
|---|---|---|---|---|
| ImageNet-100 | 1% labels | 22.50 ± 2.25<sup>*a*</sup> | 28.19 ± 1.14<sup>*a*</sup> | **28.32 ± 0.82<sup>*a*</sup>** |
|  | 10% labels | 47.05 ± 1.48<sup>*a*</sup> | **50.43 ± 1.48<sup>*a*</sup>** | 48.80 ± 0.79<sup>*a*</sup> |
| CIFAR-10 | 0.1% labels | 26.80 ± 0.76<sup>*b*</sup> | 38.76 ± 1.01<sup>*a*</sup> | **39.54 ± 0.38<sup>*a*</sup>** |
|  | 1% labels | 50.17 ± 1.94<sup>*b*</sup> | 60.12 ± 0.87<sup>*a*</sup> | **61.68 ± 0.50<sup>*a*</sup>** |
| Flowers102 | 1-shot per class | 15.46 ± 1.80<sup>*c*</sup> | 19.45 ± 1.16<sup>*b*</sup> | **25.17 ± 1.42<sup>*a*</sup>** |
|  | 5-shot per class | 39.38 ± 2.03<sup>*c*</sup> | 44.84 ± 1.96<sup>*b*</sup> | **51.65 ± 2.03<sup>*a*</sup>** |
| EMNIST Balanced | 0.1% labels | 24.62 ± 2.13<sup>*a*</sup> | 27.85 ± 2.34<sup>*a*</sup> | **29.82 ± 0.59<sup>*a*</sup>** |
|  | 1% labels | 73.05 ± 0.58<sup>*b*</sup> | 74.99 ± 0.81<sup>*a*</sup> | **75.40 ± 0.40<sup>*a*</sup>** |
| Directional GTSRB | 5-shot per class | 46.21 ± 8.59<sup>*a*</sup> | **59.81 ± 7.36<sup>*a*</sup>** | 57.16 ± 7.20<sup>*a*</sup> |
|  | 50-shot per class | 69.59 ± 7.39<sup>*a*</sup> | **89.32 ± 2.40<sup>*a*</sup>** | 83.99 ± 3.79<sup>*a*</sup> |

## Footnotes

1 clane9/imagenet-100 on the Hugging Face Hub: https://huggingface.co/datasets/clane9/imagenet-100

